# The paraventricular thalamus provides a polysynaptic brake on limbic CRF neurons to sex-dependently blunt binge alcohol drinking and avoidance behavior

**DOI:** 10.1101/2020.05.04.075051

**Authors:** Olivia B. Levine, Mary Jane Skelly, John D. Miller, Jean K. Rivera-Irizarry, Sydney A. Rowson, Jeffrey F. DiBerto, Jennifer A. Rinker, Todd E. Thiele, Thomas L. Kash, Kristen E. Pleil

## Abstract

Bed nucleus of the stria terminalis (BNST) neurons that synthesize and release the stress neuropeptide corticotropin-releasing factor (CRF) drive binge alcohol drinking and anxiety, behaviors that are primary risk factors for alcohol use disorder (AUD) and comorbid neuropsychiatric diseases more common in women than men. Here, we show that female C57BL/6J mice binge drink more than males and have greater basal BNST^CRF^ neuron excitability and synaptic excitation. We identified a dense VGLUT2+ glutamatergic synaptic input from the paraventricular thalamus (PVT) that is anatomically similar in males and females. These PVT^BNST^ neurons release glutamate directly onto BNST^CRF^ neurons but also engage a large BNST interneuron population to ultimately provide a net inhibition of BNST^CRF^ neurons, and both components of this polysynaptic PVT^VGLUT2^-BNST^CRF^ circuit are more robust in females than males. Chemogenetic inhibition of the PVT^BNST^ projection promoted binge alcohol drinking in females without affecting males, and chemogenetic activation of the pathway was sufficient to reduce avoidance behavior in both sexes in anxiogenic contexts. Lastly, we show that withdrawal from repeated binge drinking produces a female-like phenotype in the male PVT-BNST^CRF^ excitatory synapse without altering the function of PVT^BNST^ neurons *per se*. Our data describe a complex feedforward inhibitory PVT^VGLUT2^-BNST^CRF^ glutamatergic circuit that is more robust in females, plays sex-dependent roles in alcohol drinking and avoidance behavior, and undergoes sex-dependent alcohol-induced plasticity.

---

Alcohol use disorder (AUD) is highly co-expressed with other neuropsychiatric diseases including anxiety disorders, with women having increased susceptibility to this comorbidity compared to men (61% vs 35%, respectively)^1^. Binge alcohol drinking is a primary risk factor for the development of these conditions, and females across mammalian species display greater binge drinking and transition from first alcohol use to disease states more quickly than males^2^. Females also exhibit exaggerated negative health consequences across peripheral and central tissues for every drink consumed^3^. These initial differences in alcohol sensitivity suggest that the mechanisms underlying early drinking contribute to disease susceptibility and are important targets for intervention^2^. However, there is currently a lack of understanding of the mechanisms controlling binge alcohol drinking in females and sex differences in the expression of this behavior and consequent vulnerability to disease. The bed nucleus of the stria terminalis (BNST) is a hub in the brain circuits underlying anxiety and alcohol/substance use disorders in humans and is highly sexually dimorphic in mammals^4,5^. The BNST is enriched with neurons that synthesize and release corticotropin-releasing factor (CRF), a stress neuropeptide involved in the development and maintenance of anxiety and addictive disorders, and activation of BNST^CRF^ neurons drives binge drinking behavior and produces anxiety^6,7^; however, the identity and organization of upstream excitatory circuits controlling BNST^CRF^ neuron function and its role in alcohol drinking and anxiety are poorly understood. The BNST is anatomically connected to the paraventricular nucleus of the thalamus (PVT), a region likewise implicated in the etiology of alcohol and substance use disorders and associated behaviors including anxiety^8-11^. Human neuroimaging studies have demonstrated a functional connection between the thalamus and BNST that is denser in females than males^4,5^ and decreased thalamic projection strength in young individuals with alcohol abuse^12^. These converging lines of evidence implicate a sex-dependent role for the PVT-BNST projection in behavior via modulation of BNST^CRF^ neurons. Here, we systematically examined in both sexes the anatomical and functional architecture of the PVT-BNST^CRF^ circuit and its role in risky alcohol drinking and anxiety behaviors, as well as alcohol-induced plasticity.

## Results

### Greater BNST^CRF^ neuron excitation and alcohol drinking in females

First, we examined the relationship between sex-dependent binge alcohol consumption and BNST^CRF^ neuron excitation. We showed that female C57BL/6J mice consistently consume more alcohol than males using the Drinking in the Dark (DID)^13,14^ model of binge drinking (**Fig. 1a**). Using whole-cell patch-clamp slice electrophysiological recordings in BNST^CRF^ neurons (**Fig. 1b**), we found that the proportion of BNST^CRF^ neurons active in their basal state was more than twice as high in females compared to males (60% in females vs. 24% in males; **Fig. 1c**). Further evaluation of BNST^CRF^ neurons in slice showed that the frequency of spontaneous excitatory postsynaptic currents (sEPSCs), including activity-independent “miniature” EPSCs (mEPSCs), was higher in females than males, but the frequency of inhibitory postsynaptic currents (sIPSCs and mIPSCs) was not (**Fig. 1d**,**e; Supplementary Fig. 1**). These results suggest that spontaneous glutamate, but not GABA, release onto BNST^CRF^ neurons is greater in females than males, contributing to an overall increased synaptic drive onto BNST^CRF^ neurons in females (biased toward excitation) compared to males (**Fig. 1f**). To determine whether increased excitability and excitation of BNST^CRF^ neurons may be related to potentiated alcohol intake in females, we assessed the ability of chemogenetic inhibition of BNST^CRF^ neurons to suppress binge alcohol consumption in males and females. By comprehensively analyzing the effects of sex and agonist dose in a broader dataset from a mixed-sex cohort of mice we previously published^15^, we found that females with the hM4D Gi-coupled designer receptor exclusively activated by designer drug (Gi-DREADD) in BNST^CRF^ neurons required a higher dose of CNO (10 mg/kg) to attenuate binge drinking than their male counterparts (3 mg/kg; **Supplementary Fig. 2**). Together, these data suggest that the level of activity in the BNST^CRF^ neuron population was sufficiently higher in females to necessitate more robust functional inhibition to affect binge drinking behavior.

**Fig. 1:**
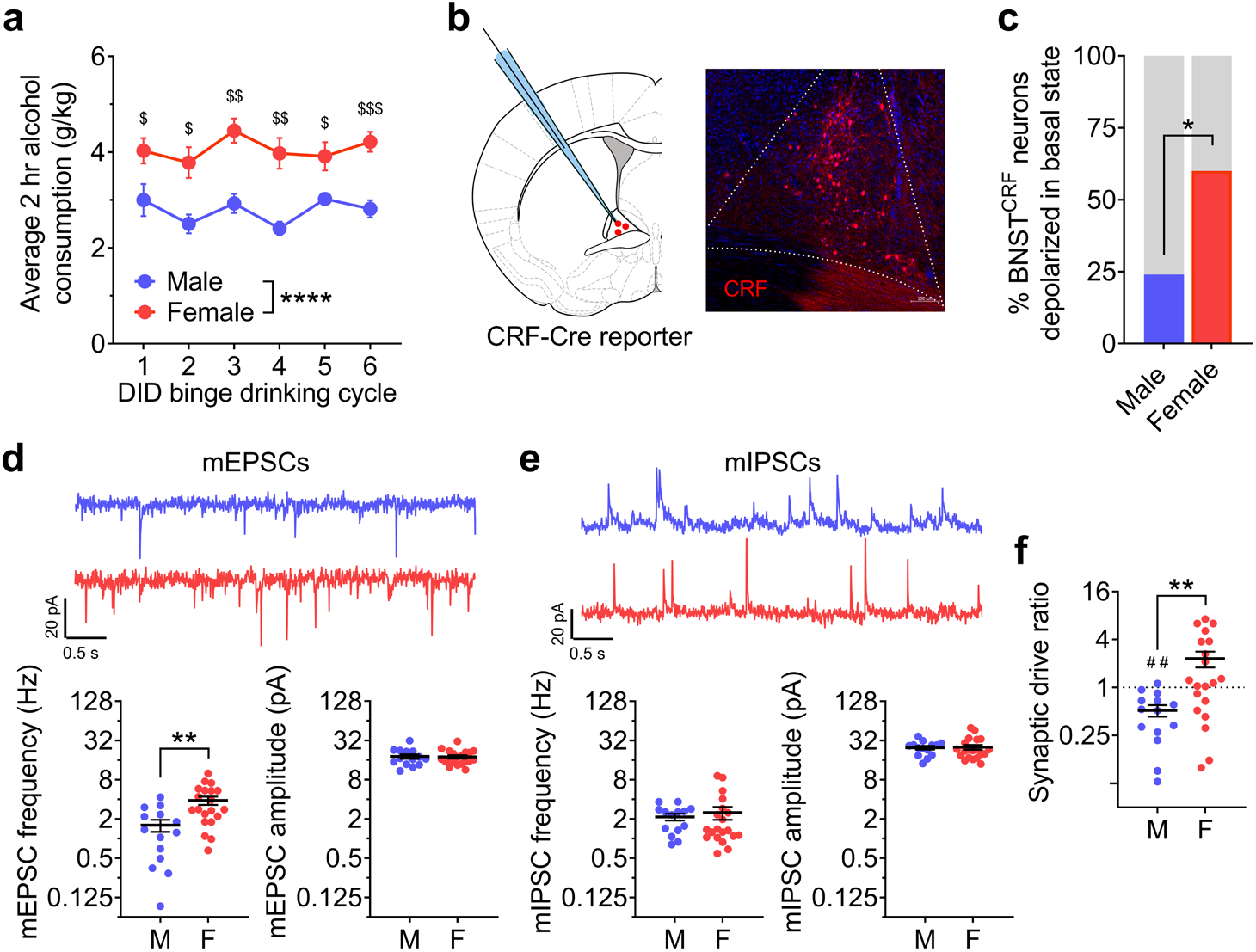
Females display higher binge alcohol drinking and have greater BNST^CRF^ neuron excitation. **a**, Average 2-hr alcohol consumption across cycles of the Drinking in the dark (DID) binge drinking paradigm showing higher binge drinking in females than males (N’s = 10 M, 10 F). 2xRM-ANOVA: main effect of sex (F_1,18_= 34.81, ^****^*P* < 0.0001) and no effect of cycle or interaction (Ps > 0.10); post hoc direct comparisons within-cycle show differences on all cycles (C1: t_17_ = 2.41, ^$^*P* = 0.032; C2: t_14.8_ = 3.41, ^$^*P* = 0.012; C3: t_17.1_ = 4.69, ^$$^*P* = 0.001; C4: t_12.4_ = 4.69, ^$$^*P* = 0.003; C5: t_11.3_ = 2.83, ^$^*P* = 0.030; C6: t_17.7_= 5.06, ^$$$^*P* = 0.0005). **b**, Schematic of whole-cell patch clamp recordings of CRF+ neurons in the BNST (BNST^CRF^ neurons, left) and representative image of a coronal BNST section from a CRF-Cre x Ai9 reporter (CRF-reporter) mouse. **c**, Proportion of BNST^CRF^ neurons sampled that are active in their basal state is greater in females than males (Fisher’s exact test ^*^*P* = 0.021, N’s = 13 M, 25 cells; 11 F, 25 cells). **d-f**, Spontaneous (miniature) excitatory and inhibitory postsynaptic currents (mEPSCs and mIPSCs) in the presence of TTX (1µM) to block action potential-dependent synaptic transmission in BNST^CRF^ neurons, (N’s = 3 M, 14 cells; 5 F, 21 cells). **d**, Top: representative traces of mEPSCs in BNST^CRF^ neurons of males (blue, above) and females (red, below). Bottom: Quantification showing that mEPSC frequency is higher in females than males (left, t_32_ = 3.44, ^**^*P* = 0.002) while mEPSC amplitude is not (right, t_32_ = 0.06, *P* = 0.956). **e**, Top: representative traces of mIPSCs in BNST^CRF^ neurons of males (blue, above) and females (red, below). mIPSC frequency and amplitude are not different between sexes (frequency, left: t_32_ = 0.32, *P* = 0.753; amplitude, right: t_32_ = 0.13, *P* = 0.901. **f**, Synaptic drive ratio, calculated as (mEPSC frequency x amplitude) / (mIPSC frequency x amplitude), in BNST^CRF^ neurons is higher in females than males (t_32_ = 2.99, ^**^*P* = 0.005) and below 1.0 in males (t_13_ = 4.06, ^##^*P* = 0.001) but not females (t_19_ = 0.83, *P* = 0.415).

### Robust projection from the paraventricular thalamus modulating BNST^CRF^ neurons

The sex differences in the underlying physiology and excitatory synaptic input to BNST^CRF^ neurons led us to examine whether there are sex differences in the anatomical and/or functional density of specific glutamatergic inputs to the BNST that may modulate BNST^CRF^ neurons. Using a viral retrograde tracing approach in VGLUT2-ires-Cre (VGLUT2-Cre) mice to label VGLUT2-positive (VGLUT2+) and negative (VGLUT2-) BNST-projecting neurons (**Fig. 2a**), we identified several known sources of excitatory input to the BNST, including a well-characterized input from the basolateral amygdala (BLA) shown to reduce avoidance behavior in males^16,17^. We found that the brain region with the densest projection to the BNST was the PVT, which was similarly robust in both sexes across the anterior-posterior extent of the PVT and enriched in the anterior-mid PVT (**Fig. 2b**,**d-e; Supplementary Fig. 3a-d**). Nearly all labeled neurons in the PVT were VGLUT2+ (**Fig. 2c; Supplementary Fig. 3e**), indicating a nearly pure glutamatergic projection, consistent with the literature showing that most PVT neurons are glutamatergic^18-21^. To determine whether PVT^VGLUT2^ neurons play a role in binge drinking behavior, we injected a Cre-dependent virus expressing the kappa opioid receptor (KOR)-based Gi-DREADD (Gi-KORD)^22^ or control (CON) virus into the PVT of VGLUT2-Cre mice (**Supplementary Fig. 4a**). Following administration of the Gi-KORD ligand Salvinorin B (SalB; 17 mg/kg, s.c.), male and female Gi-KORD mice but not CONs displayed blunted binge drinking behavior (**Supplementary Fig. 4b**). In contrast, SalB activation of the Gi-KORD did not alter sucrose drinking or avoidance behavior in the open field test (OF; **Supplementary Fig. 4c-e**), suggesting that PVT^VGLUT2^ neurons drive alcohol binge drinking behavior specifically without altering reward seeking behavior or anxiety more generally.

**Fig. 2:**
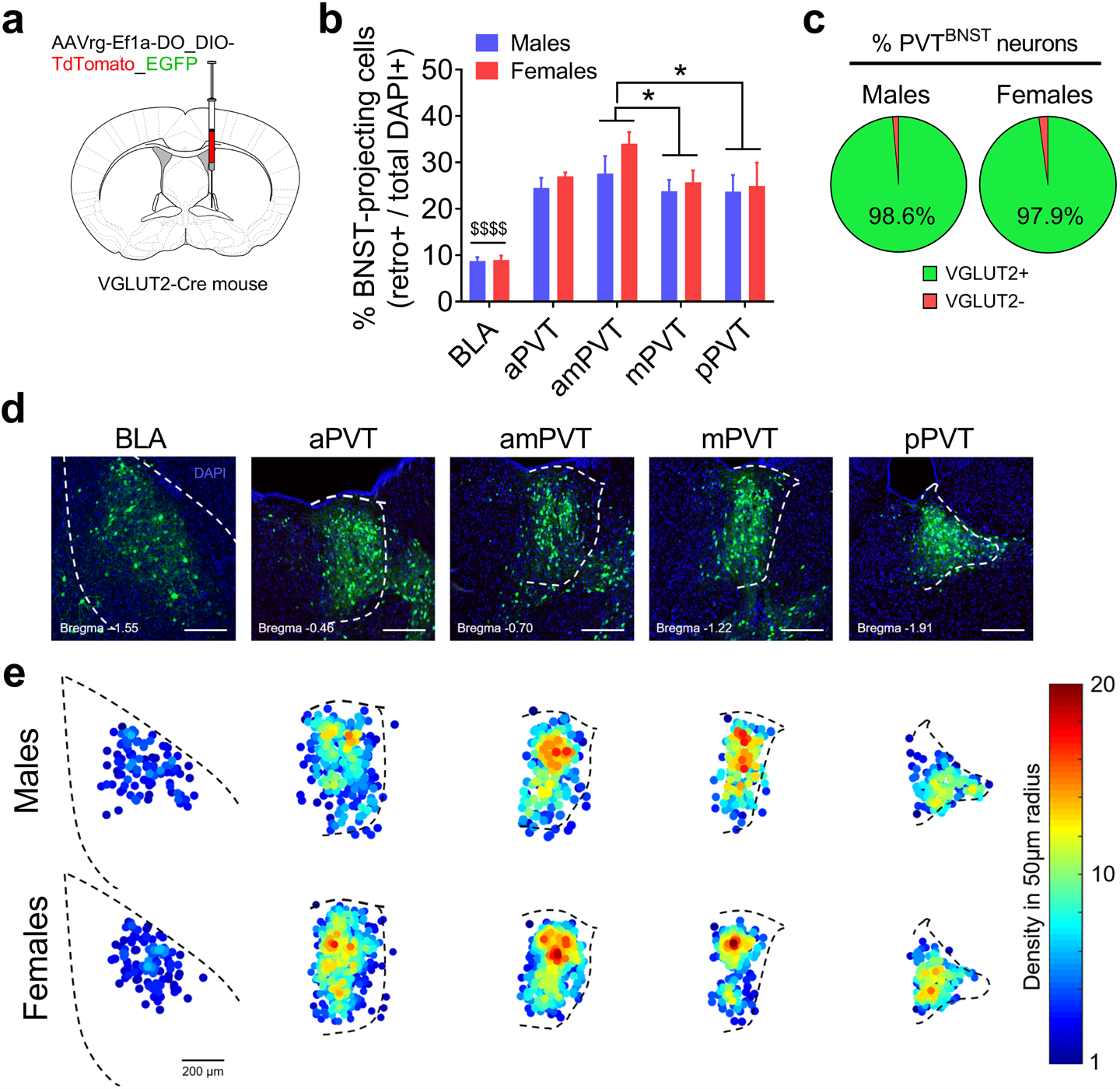
The PVT provides a dense glutamatergic projection to the BNST. **a**, Schematic of intra-BNST delivery of a virus containing a retrogradely trafficked reporter vector (AAVrg-Ef1a-DO_DIO-TdTomato_EGFP-WPRE-pA) unilaterally into the BNST of VGLUT2-Cre mice to label distal neurons that project to the BNST (N’s = 4 M, 4F). **b**, Quantification of % cells in basolateral amygdala (BLA) and paraventricular nucleus of the thalamus (PVT) sections that project to the BNST (the densest sources of glutamatergic input identified), calculated as (total # of retrogradely-labeled cells / total DAPI count) x 100. Mixed-effects model: main effect of subregion (F_4,23_ = 32.01, *P* < 0.0001, not indicated) but no effect of sex or interaction (*P*s > 0.35); post hoc direct comparisons between subregions show that the BLA has fewer BNST-projecting neurons than the PVT across all A/P coordinates (all ^$$$$^*P*s < 0.0001), and that the amPVT has more than the mPVT (t_23_ = 2.94, ^*^*P* = 0.037) and pPVT (t_23_ = 3.15, ^*^*P* = 0.027). aPVT, anterior PVT; amPVT anterior-mid PVT; mPVT, mid PVT: pPVT, posterior PVT. **c**, Proportion of VGLUT2-Cre+ (EGFP-labeled) and VGLUT2-Cre- (tdTomato-labeled) BNST-projecting PVT (PVT^BNST^) neurons in males and females, showing that almost all PVT^BNST^ neurons are VGLUT2+ in both males and females with no difference between sexes (t_6_ = 1.24, *P* = 0.261). **d**, Representative images of coronal brain slices from a virus-injected mouse illustrating the expression of DAPI (blue) and all BNST-projecting cells (both VGlut2-Cre+ and – in green) in the BLA and across the A/P extent of the PVT. **e**, Density heat maps illustrating the average number of BNST-projecting cell bodies within a 50 µM radius of each identified projection neuron for samples from each sex, matched and scaled similarly to representative images in **d**, illustrating the increased density of BNST-projecting across the PVT compared to the BLA that is similar between sexes (top: males, bottom: females).

We next evaluated sex differences in the function of BNST-projecting PVT (PVT^BNST^) neurons. Whole-cell patch-clamp recordings of PVT^BNST^ neurons indicated no sex differences in the intrinsic or synaptic excitability of this neuron population apart from decreased sIPSC amplitude in females (**Supplementary Fig. 5**). To establish whether there are sex differences in the function of the PVT-BNST (PVT^BNST^) pathway and postsynaptic responses from BNST^CRF^ neurons, we injected a CaMKIIα-driven channelrhodopsin (ChR2) virus into the PVT of CRF-reporter mice and recorded from BNST^CRF^ neurons (**Fig. 3a**,**b**). We found that ChR2 activation of the PVT synaptic input (2 ms pulse of 490 nm LED) elicited an optically-evoked monosynaptic EPSC and polysynaptic IPSC (oEPSC and oIPSC, respectively) in BNST^CRF^ neurons, as both currents could be abolished by bath application of the voltage-gated sodium channel blocker tetrodotoxin (TTX, 500 nM) but only the oEPSC was restored by addition of the voltage-gated potassium channel blocker 4-aminopyridine (4-AP, 100 μM; **Fig. 3c**,**d**). Quantification of oEPSC and oIPSC amplitudes across increasing LED power showed a more robust response in BNST^CRF^ neurons of females compared to males (**Fig. 3e**,**f, Supplementary Fig. 6a**,**b**). oIPSCs were larger than oEPSCs in most individual BNST^CRF^ neurons in both sexes, resulting in a net effect of synaptic inhibition by PVT afferent activation in both males and females (**Fig. 3g**), similar to what has been described for other limbic projections of the PVT^19-21^. In addition, the latency between optical excitation of PVT terminals and initiation of oPSCs was significantly longer for oIPSCs than oEPSCs within individual BNST^CRF^ neurons in both sexes (**Fig. 3h**), providing converging evidence that the PVT-evoked inhibition onto BNST^CRF^ neurons was polysynaptic in nature.

**Fig. 3:**
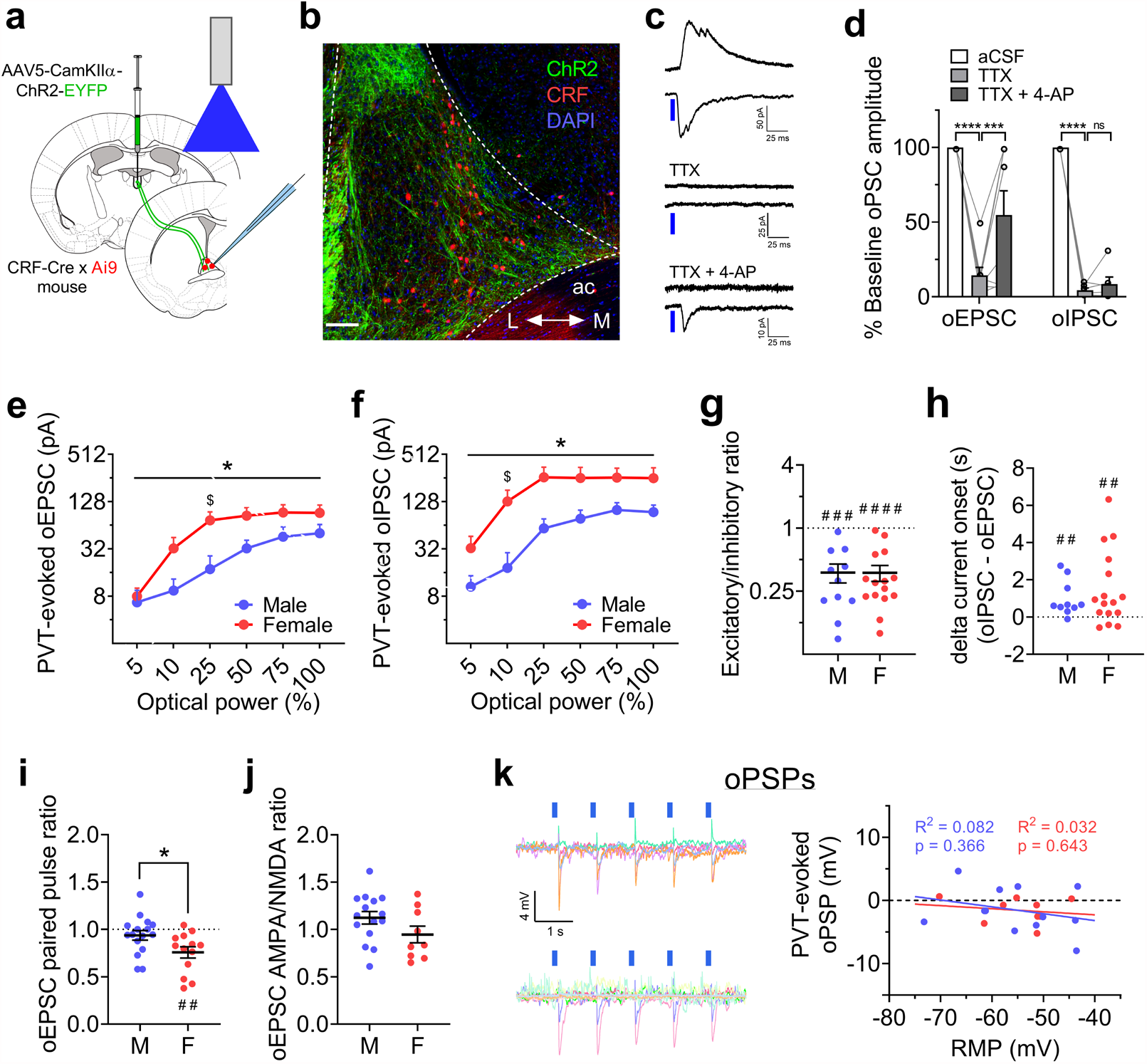
The PVT-BNST projection provides polysynaptic inhibition onto BNST^CRF^ neurons. **a-b**, Approach for slice recordings in BNST^CRF^ neurons during optical excitation of ChR2 in PVT axon terminals (**a**) and representative coronal image of the dorsal BNST (**b**); ac = anterior commissure, L = lateral, M = medial. Scale bar = 100 μM. **c-d**, Representative traces (**c**) and quantification (**d**) of time-locked PVT ChR2-evoked EPSCs and IPSCs in BNST^CRF^ neurons (oEPSCs and oIPSCs, respectively; top) in response to 2 ms pulses of blue LED, which are abolished in the presence of tetrodotoxin (TTX, 1 μM); a time-locked monosynaptic oEPSC but not oIPSC is restored with the addition of 4-aminopyridine (4-AP, 100 μM). oEPSCs: 1xRM-ANOVA effect of drug (F_2,14_= 22.6, *P* < 0.0001), post hoc t-tests (aCSF vs. TTX: t_14_ = 6.71, ^****^*P* < 0.0001; TTX vs. TTX+4AP: t_14_ = 3.16, ^**^*P* = 0.007; oIPSCs: 1xRM-ANOVA effect of drug (F_2,10_ = 429.0, *P* < 0.0001), post hoc t-tests (aCSF vs. TTX: t_10_ = 25.90, ^****^*P* < 0.0001; TTX vs. TTX+4AP: t_10_ = 1.10, *P* = 0.299). **e-f**, LED power-response curves for oEPSCs (**e**) and oIPSCs (**f**) in the same BNST^CRF^ neurons (N’s = 2 M, 6 cells; 6 F, 13 cells). **e**, 2xRM-ANOVA for oEPSCs: main effect of power (F_5,85_ = 48.74, *P* < 0.0001, not indicated) and a sex x power interaction (F_5,85_ = 2.58, ^*^*P* = 0.032), but no main effect of sex (*P* > 0.10). Post hoc direct comparisons reveal a significant difference between males and females for 25% power (t_102_ = 2.76, ^$^*P* = 0.041) but no other LED powers (*P*s > 0.55). **f**, 2xRM-ANOVA for oIPSCs: main effect of power (F_5,85_ = 54.09, *P* < 0.0001, not indicated) and a sex x power interaction (F_5,85_ = 2.89, ^*^*P* = 0.019), but no main effect of sex (*P* > 0.05). Post hoc direct comparisons reveal a significant difference between males and females for 10% power (t_102_ = 2.94, ^$^*P* = 0.024) but no other LED powers (*P*s > 0.10). **g**, oEPSC/oIPSC ratios in individual BNST^CRF^ neurons showing net synaptic inhibition in both sexes (one-sample t-tests compared to 1: males: t_10_ = 5.50,^###^*P* = 0.0001; females: t_14_ = 7.20, ^####^*P* < 0.0001) with no difference between males and females (t_24_ = 0.11, *P* = 0.912). **h**, Difference in the onset time of oIPSCs and oEPSCs. One-sample t-tests compared to 0: males: t_9_ = 3.32, ^##^*P* = 0.009; females: *t*_15_ = 3.01, ^##^*P* = 0.009, with no difference between males and females (t_24_ = 0.76, *P* = 0.455). for **g-h**, N’s = 5 M, 11 cells; 8 F, 16 cells. **i**, Paired pulse ratio of pharmacologically-isolated oEPSCs is lower in females than males (t_27_ = 2.33, \**P* = 0.028) and below 1 in females (one-sample t-test: t_12_ = 4.07, ^##^*P* = 0.002) but not males (t_15_ = 2.33, *P* = 0.235). N’s = 4 M, 16 cells; 3 F, 13 cells. **j**, No difference in oEPSC AMPA/NMDA ratios between sexes (t_22_ = 1.61, *P* = 0.121). N’s = 4 M, 15 cells; 3 F, 9 cells. **k**, Optically-evoked postsynaptic potentials in BNST^CRF^ neurons (oPSPs) from PVT ChR2+ terminals. Left: overlaid oPSP traces from basally inactive BNST^CRF^ neurons, with upward deflections indicating time-locked depolarizations and downward deflections indicating hyperpolarizations (males, top; females, bottom). Right: oPSP magnitude plotted as a function of resting membrane potential (RMP). Linear regression equations: M: Y = -0.109^*^X – 7.53; F: Y = - 0.049^$^X – 4.23. Pearson’s R^2^ values and corresponding *P* values reported in graph for males (blue) and females (red). N’s = 7 M, 12 cells; 6 F, 9 cells.

We next evaluated potential pre-vs. postsynaptic mechanisms contributing to more robust PVT-BNST^CRF^ synapses in females compared to males. While the excitability of PVT^BNST^ neurons was similar between males and female (**Supplementary Fig. 5**), paired pulse ratios of pharmacologically-isolated PVT-evoked oEPSCs in BNST^CRF^ neurons separated by 50 ms were higher in males than females, suggesting greater probability of evoked presynaptic glutamate release from PVT terminals in the BNST in females (**Fig. 3i**). In contrast, AMPA/NMDA ratios of oEPSCs were similar between males and females, indicating no postsynaptic component contributing to the sex difference in oEPSC magnitude (**Fig. 3j**). Intriguingly, PPR was below 1.0 in females (**Fig. 3i**), suggesting that these PVT terminals require relatively little excitation to elicit glutamate release and thus PVT-BNST^CRF^ synapses may serve as low-pass synaptic filters in females. In conjunction with increased spontaneous glutamate release onto BNST^CRF^ neurons in females (**Fig. 1d**), these results suggest higher PVT glutamate tone in the BNST of females. Finally, we confirmed that activating PVT^BNST^ glutamatergic inputs results in a net hyperpolarization of the membrane potential of BNST^CRF^ neurons using whole-cell current-clamp recordings. LED excitation of ChR2 from PVT terminals produced an optically-evoked postsynaptic potential (oPSP) that was negative (hyperpolarizing) in most BNST^CRF^ neurons in both males and females, regardless of their resting membrane potential (**Fig. 3k**). Evaluation of oPSPs in response to PVT ChR2 excitation across a range of frequencies (**Supplementary Fig. 6c**,**d**) showed that 20 Hz stimulation (the same frequency used to evaluate PPR) elicited a greater magnitude second oPSP than first oPSP in BNST^CRF^ neurons of males but no difference at any frequency in females, consistent with higher PPR in males than females. Thus, while female PVT inputs readily release glutamate, those in males have a lower probability of release but may be more prone to synaptic facilitation upon repeated stimulation.

### PVT^BNST^ circuit regulation of alcohol drinking and anxiety behaviors

Given the inhibitory effect of *ex vivo* PVT^BNST^ afferent activation on BNST^CRF^ neuron excitability, we next evaluated the role of the PVT^BNST^ circuit in binge drinking and anxiety-like behavior. We used a multiplexed chemogenetic strategy^22^ to bidirectionally and independently manipulate the PVT^BNST^ circuit during behavior by injecting either a cocktail of a Cre-dependent excitatory (hM3D) Gq-DREADD + Cre-dependent inhibitory Gi-KORD or a Cre-dependent CON vector in retrogradely Cre-labeled PVT^BNST^ neurons (**Fig. 4a**,**b; Supplementary Fig. 7**). We found that CNO administration (5 mg/kg, i.p.) to activate the Gq-DREADD prior to alcohol access during Day 4 DID did not reliably alter binge consumption of 20% alcohol compared to vehicle baseline (**Fig. 4c**); however, there was trend for a decrease in consumption in DREADD males (see **Supplementary Fig. 8a**), suggesting that activation of the PVT^BNST^ pathway may be sufficient to blunt binge alcohol drinking in a subset of males. In contrast, Salvinorin B (SalB) administration (17 mg/kg, s.c.) to activate the Gi-KORD robustly increased binge alcohol drinking in DREADD females but had no effect in DREADD males or either CON group (**Fig. 4d, Supplementary Fig. 8b**), suggesting that tonic activity of the PVT^BNST^ circuit (**Supplementary Fig. 5c**) engaging interneurons is necessary for active suppression of alcohol drinking behavior in females. As BNST^CRF^ neurons show larger postsynaptic responses to the PVT^BNST^ input in females, reducing the activity of the PVT^BNST^ input may result in a more robust disinhibition of BNST^CRF^ neurons to further enhance binge drinking. Intriguingly, neither chemogenetic manipulation affected 1% sucrose consumption in a similar DID paradigm (**Supplementary Fig. 8c**,**d**). These results suggest that the PVT^BNST^ projection plays a unique inhibitory role in binge drinking behavior that is not dependent on modulation of the general rewarding or aversive aspects of drug intake and consummatory behavior previously shown to be mediated by other major limbic outputs of the PVT such as the nucleus accumbens^20,23,24^.

**Fig. 4:**
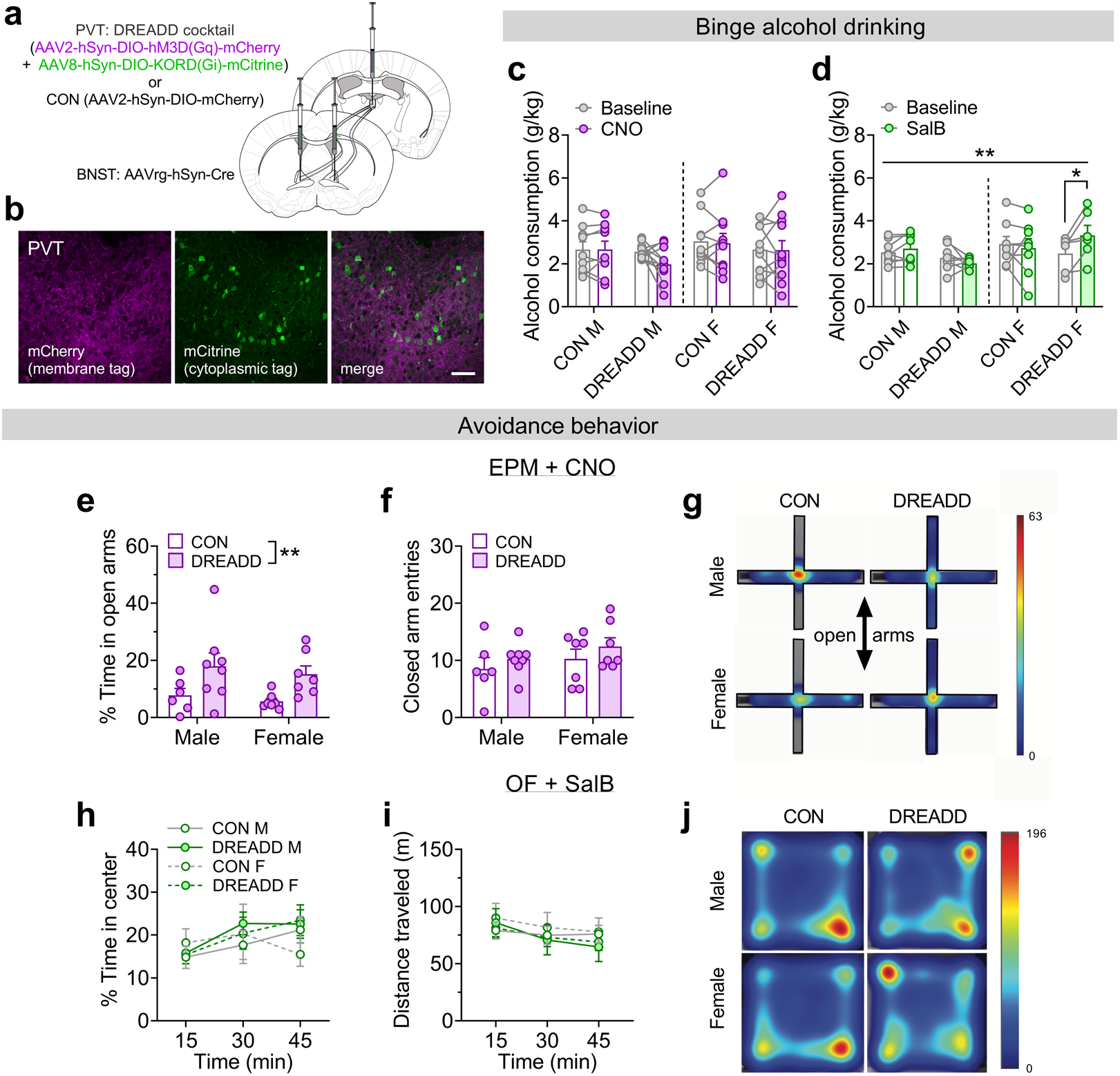
The PVT-BNST projection sex-dependently regulates binge alcohol drinking and avoidance behavior. **a**, Viral strategy for a multiplexed chemogenetic approach to bidirectionally manipulate the PVT-BNST (PVT^BNST^) projection during behavior. **b**, Representative image of the PVT showing coexpression of the Gq-DREADD (membrane mCherry fusion protein in purple) and Gi-KORD (separated cytoplasmic mCitrine tag in green) in PVT^BNST^ neurons. Scale bar = 50 μM. **c-d**, Two-hr alcohol consumption during within-DID cycle vehicle baseline vs. Day 4 ligand administration in CON and DREADD mice. **c**, CNO administration (5 mg/kg, i.p., Gq-DREADD activation). 3xRM-ANOVA: no effects of sex, DREADD, CNO, or any interactions between the variables (*P*s > 0.15, but see **Supplementary Fig. 6a** for further analysis; N’s = 9 CON M, 11 DREADD M, 10 CON F, 11 DREADD F). **d**, SalB administration (17 mg/kg, s.c., Gi-KORD activation). 3xRM-ANOVA: sex x DREADD x SalB interaction (F_1,26_ = 8.76, ^**^*P* = 0.007) and no main effects and no interactions between two variables (*P*s > 0.05); post hoc paired t-tests: SalB increased alcohol consumption in DREADD females (t_26_ = 3.08,^*^*P* = 0.019) but not DREADD males or either control group (*P*s > 0.65). N’s = 8 CON M, 8 DREADD M, 8 CON F, 6 DREADD F, with 0-3 mice excluded per group due to baseline drinking below criteria. **e-g**, Effects of CNO (Gq-DREADD activation) on behavior in the elevated plus maze (EPM). N’s = 6 CON M, 8 DREADD M, 7 CON F, 7 DREADD F. **e**, Percent time spent on the open arms is higher in DREADD mice. 2xANOVA: main effect of DREADD (F_1,24_ = 9.30, ^**^*P* = 0.006) but no other effects (*P*s > 0.45). Post hoc direct comparisons within sex show no significant differences (*P*s > 0.05). **f**, Number of closed arm entries is not different. 2xANOVA: no effects or interaction between sex and DREADD (*P*s > 0.20). **g**, Representative tracking heat maps. **h-j**, No effect of SalB (Gi-KORD activation) on behavior in the open field test (OF). N’s = 4 CON M, 5 DREADD M, 4 CON F, 5 DREADD F. **h**, Percent time spent in center. 3xRM-ANOVA: main effect of time (F_2,27.6_ = 8.70, *P* = 0.003, not indicated) and no other effects or interactions (*P*s > 0.10). **i**, Distance traveled. 3xRM-ANOVA: main effect of time (F_1.2,17.4_ = 16.1, *P* = 0.0005, not indicated) but no other effects or interactions (*P*s > 0.10). **j**, Representative tracking heat maps.

We also probed whether the PVT^BNST^ circuit plays a role in anxiety-like behavior, given our previous work showing that activity of the BNST^CRF^ neuron population is anxiogenic^15,25^ and that the PVT regulates emotional behaviors^11,19^. CNO administration to activate the Gq-DREADD decreased avoidance of the open arms on the elevated plus maze (EPM) in DREADD mice compared to CON mice (**Fig. 4e-g**), an anxiolytic effect of Gq-DREADD activation that we replicated in the open field test (OF; **Supplementary Fig. 8e**,**f**). In contrast, SalB administration to activate the Gi-KORD did not affect avoidance of the center of the open field (**Fig. 4h-j**). Together, these results suggest that in an anxiogenic context when BNST^CRF^ neuron activity is high^26,27^, activation of the PVT^BNST^ synaptic brake is sufficient to decrease anxiety-like behavior.

### Repeated alcohol use produces a female-like phenotype in BNST^CRF^ synapses

Finally, we examined the sex-dependent plasticity in this circuit following repeated binge alcohol drinking to understand whether there are changes in the PVT^VGLUT2^-BNST^CRF^ circuit during withdrawal that could precipitate or contribute to increased disease vulnerability. We found that one day after three cycles of EtOH DID (**Fig. 5a**), the proportion of BNST^CRF^ neurons in an active state was increased following alcohol exposure in males (**Fig. 5b**); intriguingly, this proportion was unchanged by a history of alcohol exposure in females, suggesting that basal BNST^CRF^ neuron population-level activity may be near a maximum in naÏve females (**Fig. 1b; Fig. 5b**) in the absence of a discrete salient stimulus (such as during an anxiety assay such as EPM here).

**Fig. 5:**
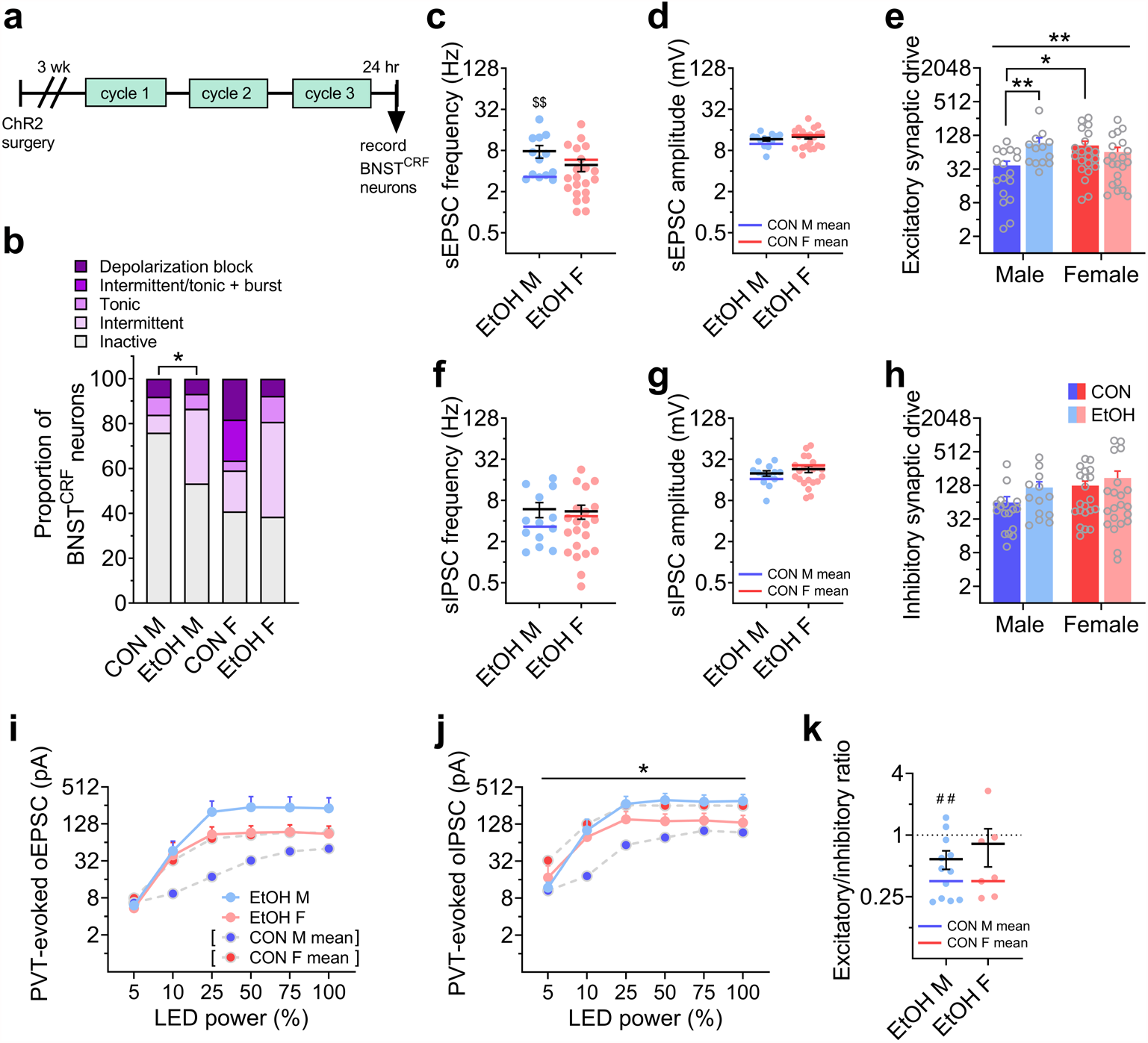
A history of voluntary binge alcohol drinking induces a female-like phenotype in male BNST^CRF^ neurons and PVT-BNST^CRF^ synapses. **a**, Experimental timeline for recording from BNST^CRF^ neurons one day following three cycles of alcohol DID (EtOH) or a water DID control (CON) procedure (means for CON cell data presented in **Figs. 1 & 3** are represented in this figure by blue and red lines for CON M and F, respectively). **b**, % BNST^CRF^ neurons in various states of excitability. Alcohol drinking increases the proportion of depolarized neurons in males (Fisher’s exact test \**P* = 0.037) but not females (*P* > 0.99). N’s = 8 EtOH M, 23 cells; 10 EtOH F, 26 cells. **c-h**, Effect of repeated binge alcohol drinking on spontaneous EPSCs and IPSCs in BNST^CRF^ neurons (N’s = 5 EtOH M, 13 cells; 8 EtOH F, 21 cells). **c**, sEPSC frequency. Males: t_28_ = 3.08, ^$$^*P* = 0.005; females: t_40_ = 1.02, *P* = 0.314). **d**, sEPSC amplitude. Males: t_28_ = 1.55, *P* = 0.132; females: t_40_ = 0.520, *P* = 0.606. **e**, Excitatory synaptic drive, calculated as sEPSC frequency x sEPSC amplitude. 2xANOVA: sex x EtOH interaction (F_1,68_ = 10.55, ^**^*P* = 0.002) but no main effects (*P*s > 0.05); post hoc direct comparisons show a sex difference between CON M and F (t_68_ = 3.10, ^*^*P* = 0.014) and effect of EtOH in males (t_68_ = 3.37, ^$$^*P* = 0.007) but not females (*P* > 0.50). **f**, sIPSC frequency. Males: t_28_ = 1.48, *P* = 0.151; females: t_40_ = 0.02, *P* = 0.981. **g**, sIPSC amplitude. Males: t_28_ = 1.16, *P* = 0.257; females: t_40_ = 1.02, *P* = 0.314. **h**, Inhibitory synaptic drive. 2xANOVA: no effects or interactions (*P*s > 0.30). **i-j**, Power-response curves for PVT-evoked oEPSCs (**i**) an oIPSCs (**j**). N’s = 4 EtOH M, 12 cells and 3 EtOH F, 8 cells. **i**, 3xRM-ANOVA on oEPSCs: main effect of power (F_5,175_ = 86.4, *P* < 0.0001, not indicated) and a trend for an EtOH x power interaction (F_5,175_ = 2.25, *P* = 0.052), but no other effects or interactions (*P*s > 0.15); see **Supplementary Fig. 12** for further analysis. **j**, 3xRM-ANOVA on oIPSCs: main effect of power (F_5,175_ = 103.4, *P* < 0.0001, not indicated) and a sex x EtOH interaction (F_1,35_ = 4.71, ^*^*P* = 0.037), but no other effects or interactions (*P*s > 0.05). **k**, oEPSC/oIPSC ratio in BNST^CRF^ neurons of EtOH mice. EtOH M have E/I ratios below 1 (one-sample t-test: t_12_ = 3.28, ^##^*P* = 0.007), which does not differ from that of CON M (t_21_ = 1.55, *P* = 0.137). EtOH F do not have an E/I ratio below 1.0 (one-sample t-test: t_6_ = 1.71, *P* = 0.137), however this ratio does not significantly differ from CON F (t_20_ = 1.84, *P* = 0.081).

Alternatively, the population level excitability of BNST^CRF^ neurons in females may be relatively impervious to the effects of extended alcohol exposure. Notably, a range of neural activity patterns were observed in all groups, illustrating the heterogeneity of the BNST^CRF^ population in both sexes as previously described in males^28,29^. Most other measures of intrinsic excitability and current-injected firing were neither different between males and females nor altered following EtOH DID exposure when measured either at their resting membrane potential (RMP; **Supplementary Fig. 9**) or at a common hyperpolarized potential of -70 mV (**Supplementary Fig. 10**). This suggests that the primary effect of repeated alcohol exposure on BNST^CRF^ neuron excitability is on the population-level activity, with males developing a female-like phenotype. Examination of synaptic transmission showed that sEPSC frequency was increased in EtOH males but unchanged in females (**Fig. 5c**), while sEPSC amplitude was unchanged in both sexes (**Fig. 5d**). This led to an increase in excitatory synaptic drive onto BNST^CRF^ neurons in males but not females (**Fig. 5e**). In contrast, there were no changes in the frequency, amplitude, or synaptic drive of sIPSCs in either sex (**Fig 5f-h**). Assessment of the kinetics of sPSCs revealed increased half-width and weighted tau of sEPSCs in EtOH males, consistent with an increase in glutamatergic transmission (**Supplementary Fig. 11**).

We further examined plasticity in PVT-evoked postsynaptic responses from BNST^CRF^ neurons and found no significant effect of EtOH on oEPSC raw amplitude across increasing LED power (**Fig. 5i**). However, when oEPSC amplitude was normalized to maximum within each cell, there was a leftward shift in the power-response curve in males but not females (**Supplementary Fig. 12a**,**b**), demonstrating that lower optical power was able to elicit larger oEPSCs in EtOH males compared to controls. Further, analysis of the slope of the initial increase in raw oEPSC amplitude across 5-25% LED power confirmed this alcohol-induced increase in the excitatory postsynaptic responses of BNST^CRF^ neurons in males (**Supplementary Fig. 12c**,**d**), mirroring the alcohol exposure effect on sEPSCs in males (**Fig. 5c**) and suggesting that PVT-BNST^CRF^ synapses comprise a large contingent of the glutamate synapses vulnerable to alcohol-induced plasticity. oIPSCs were also somewhat enhanced by alcohol exposure, particularly in males, as analysis of raw oIPSCs across LED power showed a sex x EtOH interaction (**Fig. 5j**), however normalized oIPSC amplitude was not significantly altered in BNST^CRF^ neurons of EtOH mice in either sex (**Supplementary Fig. 12e**,**f**); initial slope was increased in EtOH mice, but this was not significantly different specifically within either sex (**Supplementary Fig. 12g**,**h**). Following voluntary alcohol exposure, the E/I ratio of oPSCs remained significantly below 1.0 in males but not females (**Fig. 5k**), suggesting that while the overall E/I balance toward inhibition was maintained in males (that is, increased PVT-mediated excitation was met with increased polysynaptic inhibition), the net inhibitory effect of PVT afferent activation at BNST^CRF^ neurons was attenuated in females. However, oEPSC/oIPSC ratios were not different between CON and EtOH females, suggesting that this statistical lack of E/I ratio below one may be due to increased variability in the responses of individual neurons/synapses in females rather than a robust loss of polysynaptic inhibition, Notably, we observed very few sex differences or alcohol-induced changes in the electrophysiological properties, excitability, and synaptic transmission of PVT^BNST^ neurons themselves (**Supplementary Figs. 13 and 14**), suggesting that sex- and alcohol-dependent differences in the PVT-BNST^CRF^ synapse strength are not due to differences in PVT^BNST^ neuronal excitability *per se*. Together, our data suggest that the effects of sex and repeated binge alcohol drinking are on the tonic activity of the BNST^CRF^ neuron population, due to basal differences in and differential plasticity in PVT^VGLUT2^-BNST^CRF^ synapses and BNST^CRF^ neuron responses to the largely tonically-active PVT^BNST^ afferents.

## Discussion

Altogether, we found that female mice display more robust binge drinking behavior than males and that BNST^CRF^ neurons are both more tonically active and receive increased excitatory synaptic input in females relative to males. We further demonstrated that the PVT provides a dense excitatory input to the BNST in both sexes^30^ that is functionally more robust in females than males, consistent with human neuroimaging studies on thalamic outputs^4,5^. We report that while PVT glutamate neurons directly synapse onto BNST^CRF^ neurons as recently shown anatomically in males^30^, they also provide robust feedforward inhibition of this population by recruiting interneurons, providing a synaptic “brake” on BNST^CRF^ neuron excitability; such feedforward inhibition has also been reported for other PVT projections, such as those to the nucleus accumbens (NAc) and central amygdala (CeA)^20,21^, to modulate appetitive, drug, and fear-related behaviors in males^19-21,24,31^. For example, the PVT regulates appetitive learning and behavioral responses for availability of sucrose, food, and water, particularly through its projection to the NAc^32-34^. Intriguingly, we found that inhibition of the entire PVT glutamate neuron population suppressed voluntary, predictable binge drinking behavior (without an effect on palatable binge sucrose consumption using a parallel paradigm; Supplementary Fig. 4), consistent with many of these previously reported complex roles of the PVT in motivated behaviors. In contrast, we found that the PVT^BNST^ circuit plays a sex-dependent but opposite role in binge alcohol drinking, such that removal of the PVT^BNST^ brake disinhibits alcohol consumption in females without affecting sucrose intake. In contrast, PVT^BNST^ activation may inhibit alcohol consumption in a subset of males, pointing to a difference in the set point for engagement of the PVT-BNST^CRF^ circuit (and thus modulation) between sexes. Our findings are some of the first to provide insight into the mechanisms underlying binge alcohol drinking in females. They also highlight underlying circuit features that are different between males and females, conferring complex sex-dependent control and circuit modulation of binge alcohol consumption. They further point to a unique role for the PVT^BNST^ circuit in alcohol drinking that is independent of the role of the PVT in general appetitive reward.

In addition, we found that activation of the PVT^BNST^ pathway in aversive contexts, when BNST^CRF^ neuron activity is high^26,27^, is sufficient to reduce avoidance behavior in both sexes, similar to the role of the BLA input in males^16^. These results suggest a critical sex-dependent role of this thalamo-limbic circuit in the control of and relationship between risky alcohol drinking and anxiety states. Further, repeated binge drinking produced a female-like phenotype in PVT-BNST^CRF^ synapses and BNST^CRF^ neurons in males, characterized by increased excitation observed in naïve females and associated with greater binge drinking behavior. Notably, while the overall excitability phenotype becomes more similar to a female state, the mechanism(s) driving the alcohol-induced increase in excitability in males may be distinct from the underlying mechanisms of sex differences in excitability in the naïve population. However, our results provide a solid basis for future examination of the molecular similarities and differences between BNST^CRF^ neurons in males and females, as well as the effects of chronic alcohol. Given the relationship between high activity of this population and the associated behavioral phenotypes (which could include additional sex-dependent behavioral phenotypes), this circuit may be a target for sex-specific interventions in individuals with risky binge drinking behaviors and resulting psychiatric diseases including AUD and anxiety disorders. In addition, other glutamatergic inputs and the BNST^CRF^ neuron population itself interact highly with other BNST neuron subpopulations, including PKCδ^30^, neuropeptide Y^15^, and dynorphin^17^, among others. However, the literature is not yet comprehensive and provides some directly conflicting evidence regarding the independence and interactions between some BNST subpopulations. For example, one recent report suggests that the CRF and PKCδ populations in the male BNST receive direct synaptic input from the PVT and other regions as independent subpopulations that are inhibitory upon one another and modulate anxiety-like behavior in opposing manners (with CRF neurons being anxiogenic)^30^. In contrast, another recent study shows that these are highly overlapping populations and that acute stress recruits additional PKCδ expression in CRF neurons in females but not males^35^, suggesting an important and perhaps sex-dependent function of the overlapping population of neurons. While both studies find that stress activates both types of neurons, they come to different conclusions about the organization and role(s) of these neurons in behavior. Further, BNST^CRF^ neurons are themselves complex, serving as both projection and interneurons^25^; as such, some may laterally inhibit others to participate in the polysynaptic inhibition we observed here. Altogether, our results and others point to the need for future studies to disentangle the independence, overlap, and interactions between BNST subpopulations, with special attention paid to the effects of sex and various physiological stressors (including alcohol and other drugs of abuse). Understanding the microcircuit organization and molecular makeup of these neurons is critical to defining the broader circuit architecture containing the PVT^VGLUT2^-BNST^CRF^ circuit described here.

## Methods

### Subjects

All experimental mice were male and female adult mice on a C57BL/6J background strain. Wild-type C57BL/6J mice were purchased as adults from Jackson Laboratory, and all transgenic lines were bred in our animal facility. CRF-ires-Cre (CRF-Cre)^15,36^ and VGLUT2-ires-Cre (VGLUT2-Cre)^37^ mice were bred with WT C57BL/6J mice, and hemizygous CRF-Cre mice were bred with homozygous floxed Ai9-tdTomato or floxed EGFP-L10a mice purchased from Jackson Laboratory (stocks 007909 and 024750) to produce CRF-Cre-reporter mice. Mice were group housed with *ad libitum* access to food and water in colony room on a 12:12 hr reverse light cycle, with lights off at 7:30 a.m. Mice were singly housed for one week prior to the onset of behavioral experiments and remained singly housed thereafter. Experiments began approximately 3 hr into the dark phase of the light cycle. All experimental procedures were approved by the Institutional Animal Care and Use Committees at Weill Cornell Medicine and University of North Carolina-Chapel Hill.

### Behavior assays

The standard Drinking in the Dark (DID) binge alcohol drinking paradigm was used in mice to model human binge consumption behavior^14^. For each cycle of EtOH DID, three hr into the dark cycle, the home cage water bottle was replaced with a bottle containing 20% (v/v) alcohol (EtOH) for two hr on Days 1-3 and four hr on Day 4, followed by three days of forced abstinence between consecutive cycles. A similar access schedule was used to evaluate binge sucrose consumption, except that home cage water bottles were replaced with 1% (w/v) sucrose. For all drinking experiments, empty “dummy” cages on the same rack as behavior mice received the same EtOH or sucrose bottle replacement, and consumption was adjusted for leak from dummy bottles and then normalized to bodyweight. The open field test (OF) was used to evaluate avoidance and locomotor behavior as previously described^15^. Mice were placed in the 50 x 50 cm arena for 60 min, and Ethovision video tracking (Noldus, Wageningen, Netherlands) was used to quantify raw locomotor and location data used to calculate measures including distance traveled and time spent in each compartment (center vs. periphery, total). The elevated plus maze (EPM) was also used to assess anxiety-like behaviors and was conducted in a plexiglass maze with two open and two closed arms (35 cm length x 5.5 cm width, with walls 15 cm tall over the closed arms). Mice were placed in the center of the EPM for five-minute trials and movement and time spent in each compartment was tracked using Ethovision. Total time and percent time spent in each arm were quantified.

### Stereotaxic surgeries

For experiments requiring site-directed administration of viral vectors or retrobeads, mice were anesthetized with 2% isoflurane (VetEquip, Livermore, CA) in 0.8% oxygen in an induction chamber (VetEquip, Livermore, CA) then placed in an Angle Two mouse stereotaxic frame (Leica Biosystems, Wetzlar, Germany) and secured with ear bars into a nose cone delivering isoflurane to maintain anesthesia. Mice were given a subcutaneous injection of meloxicam (2 mg/kg) for preemptive analgesia and 0.1 mL of 0.25% Marcaine around the incision site. A Neuros 7000 series 1 µL Hamilton syringe with 33-gauge needle (Reno, NV) connected to a remote automated microinfusion pump (KD Scientific, Holliston, MA) was used for construct delivery at a rate of 50-100 nL/min to the PVT (A/P: -0.82, M/L: 0.00, D/V: -3.25, 200 nL) or BNST (A/P: +0.3 mm, M/L: ±1.1 mm, D/V: -4.35 mm, 250 nL). Following infusion, the needle was left in place for 10 min and then slowly manually retracted to allow for diffusion and prevent backflow of virus. Mice were continuously monitored for at least 30 minutes post-surgery to ensure recovery of normal breathing pattern and sternal recumbency, and then checked daily.

### in vivo chemogenetic manipulations

To examine the role of PVT^VGLUT2^ neurons in behavior, VGLUT2-Cre mice received a midline stereotaxic injection into the PVT of the kappa opioid receptor Gi-coupled DREADD (Gi-KORD) AAV9-hSyn-DIO-HA-KORD-IRES-mCitrine (gift from Bryan Roth) or control vector AAV8-hSyn-DIO-EYFP into the PVT (200 nL; Supplementary Fig. 4a). To examine the role of the PVT^BNST^ projection in binge alcohol drinking, binge sucrose drinking, and avoidance behavior, we used a multiplexed DREADD approach. C57BL/6J mice received injections of a retrograde Cre virus (AAVrg-hSyn-HI.EGFP-Cre.WPRE.SV40 or AAVrg-pmSyn1-EBFP-Cre) bilaterally in the BNST a midline PVT injection of either: a) 1:1 cocktail of the excitatory Gq-DREADD AAV2-hSyn-DIO-hM3D(Gq)-mCherry (125 nL) plus inhibitory Gi-KORD AAV8-hSyn-DIO-KORD(Gi)-mCitrine (125 nL) or b) control virus AAV2-hSyn-DIO-mCherry (250 nL, **Fig. 4a**,**b**). Approximately one week following surgery, mice started the DID procedure with a baseline cycle followed by a cycle in which they received 0.9% sterile saline vehicle (10 ml/kg, i.p.) injections on Days 2 and 4 forty min prior to alcohol access to habituate to the injection procedure. Chemogenetic manipulations began the next cycle with saline on Day 2 and clozapine-n-oxide (CNO, 5 mg/kg in 0.9% saline) on Day 4. The KORD was subsequently evaluated similarly with a DMSO vehicle (1 ml/kg, s.c.) injection cycle and manipulation cycle with Salvinorin B (SalB, 17 mg/kg in DMSO). Mice displaying injection stress defined by > 1g/kg reduction in and < 1g/kg vehicle baseline drinking were excluded from analysis. For the OF and EPM, half of the animals of each sex received vehicle and half received the DREADD activator (CNO or SalB) 40 min prior to the assay. For sucrose DID, the same SalB or CNO drug administration procedure was used as that for EtOH DID.

### Brain extraction and fluorescence immunohistochemistry

Following behavior procedures, mice were deeply anesthetized with pentobarbital (100 mg/kg, i.p.) and transcardially perfused with sterile phosphate-buffered saline (PBS) followed by 4% paraformaldehyde (PFA). Brains were extracted, post-fixed overnight in 4% PFA, and then placed in PBS until they were sliced on the coronal plane in 45 μm sections on a VT1000S vibratome (Leica Biosystems) to check injection placements and viral expression (hit maps of these expression data are presented in **Supplementary Fig. 7a**,**b**). To amplify the expression of fluorophore tags, coronal slices containing the PVT and BNST from DREADD and CON mouse brains underwent immunofluorescence staining. Slices were washed twice in PBS followed by 0.2% triton (Fisher Bioreagents, Hampton, NH) for 10 minutes each and then blocked in 5% normal donkey serum (NDS) for 30 minutes. Tissue was incubated in primary antibody (DsRed rabbit polyclonal 1:400, Takara, Kusatsu, Japan) overnight at room temperature. The next day, slices were washed 3 times in 0.2% triton for 10 minutes each and then blocked in 5% NDS for 30 minutes. Tissue was then incubated in secondary antibody (Alexafluor-568 donkey anti-rabbit, 1:250, Invitrogen, Carlsbad, CA) for 2 hours and subsequently washed twice in 0.2% Triton and then in PBS for 10 minutes each. Slices were counterstained with DAPI, and mounted on slices and coverslipped with Vectashield hardmount antifade mounting medium (Vector Labs, Burlingame, CA) and stored in the dark at 4°C until imaged (as described below) to verify surgical placements and viral expression.

### Retrograde neuronal tracing

For anatomical circuit tracing of excitatory inputs to the BNST, VGLUT2-Cre mice received stereotaxic injections of a retrograde virus (AAVrg-Ef1a-DO-DIO-TdTomato_EGFP-WPRE-pA) unilaterally into the BNST, which was is retrogradely trafficked to ultimately express GFP in Cre+ cells and tdTomato in Cre-cells that project to the BNST. Following three weeks to allow for optimal viral expression, mice were sacrificed and their brains harvested for quantification as described below.

### Image acquisition and analysis

Coronal brain slices were collected and imaged for all experiments to confirm and quantify viral expression and immunolabeling. Images were acquired on a Zeiss LSM 880 Laser Scanning Confocal microscope (Carl Zeiss, Oberkochen, Germany). For the retrograde tracing experiment, images of the PVT and BLA were quantified using ImageJ (US National Institute of Health) to count the total number of DAPI-stained nuclei, GFP+ and tdTomato+ cells. The coordinates of each cell were analyzed using custom a MATLAB (MathWorks, Natick, MA) program and normalized to the most dorsomedial point of the PVT. Heatmaps of the density of BNST-projecting neurons were generated as described elsewhere^38^, overlaying data from four mice per sex. The proportion of PVT projectors that are VGLUT2+ was calculated as GFP+ / (GFP+ plus tdTomato+). Coordinates analyzed (mm from Bregma): BLA (−1.55), anterior (aPVT, -0.46mm), anterior-mid (am-PVT, - 0.70), mid (mPVT, -1.22), and posterior (pPVT, -1.91).

### ex vivo slice electrophysiology and calcium imaging

Slice electrophysiology experiments were performed as previously described^6,15^. Mice were decapitated under isoflurane anesthesia and their brains rapidly extracted. Coronal BNST slices (300 µm) were prepared on a VT1200 vibratome (Leica Biosystems) in ice-cold, oxygenated (95% O_2_/5% CO_2_) sucrose artificial cerebrospinal fluid (aCSF) containing (in mM): 194 sucrose, 20 NaCl, 4.4 KCl, 2 CaCl2, 1 Mg Cl2, 1.2 NaH2PO4, 10 glucose, and 26 NaHCO3 (pH 7.3 and 30 mOsm). Slices were transferred to a holding chamber with 30 °C oxygenated normal aCSF (in mM):124 NaCl, 4.4 KCl, 2 CaCl2, 1.2 MgSO4, 1 NaH2PO4, 10 glucose, and 26 NaHCO3 (pH 7.3 and 30 mOsm) and allowed to equilibrate for at least one hour. For electrophysiological recordings, slices were transferred to a submerged recording chamber (Warner Instruments, Hamden, CT) and perfused at a rate of 2 mL/min with 30°C oxygenated normal aCSF.

BNST^CRF^ neurons were identified for recording with their tdTomato tag using a 580 nm LED under 40x objective (Olympus, Tokyo, Japan). Signals were acquired using a Multiclamp 700B amplifier (Molecular Devices), digitized, and analyzed via pClamp 10 or 11 software (Molecular Devices). Input resistance and access resistance were continuously monitored throughout experiments, and cells in which access resistance changed by more than 20% were not included in data analysis. Excitability experiments were performed in current-clamp configuration using a potassium gluconate-based intracellular recording solution containing (in mM): 135 KGluc, 5 NaCl, 2 MgCl_2_-6H2O, 10 HEPES, 0.6 EGTA, 4 Na-ATP and 0.4 Na-GTP (pH 7.3 and 290 mOsm). Synaptic transmission was measured in voltage-clamp configuration using a cesium-methanesulfonate-based intracellular recording solution containing (in mM): 135 CsMeth, 10 KCl, 10 HEPES, 1 MgCl_2_·6H2O, 0.2 EGTA, 4 Mg-ATP, 0.3 Na_2_GTP, 20 phosphocreatine (pH 7.3, 290 mOsm) and driving force (−55 mV and +10 mV) to isolate excitatory and inhibitory transmission within individual neurons. Miniature postsynaptic currents were measured in the presence of tetrodotoxin (TTX, 1 µM) in the aCSF bath. Paired pulse ratio and AMPA/NMDA ratio were assessed using a cesium gluconate-based intracellular recording solution containing (in mM): 117 D-gluconic acid, 20 HEPES, 0.4 EGTA, 5 TEA, 2 MgCl_2_-6H2O, 4 Na-ATP and 0.4 Na-GTP (pH 7.3 and 290 mOsm), at a holding potential of -70 mV, and with picrotoxin (25 µM) in the aCSF bath.

Prior to slice electrophysiology experiments, mice underwent three cycles of EtOH DID (EtOH group) or a water control DID procedure in which the replacement bottle contained water instead of EtOH (CON group) to allow for investigation of basic sex differences in function as well as for direct comparison to EtOH mice. Twenty-four hr following the onset of the last EtOH or water bottle access, mice were sacrificed for slice electrophysiology experiments as described above. For experiments investigating PVT^BNST^ neurons, mice received bilateral intra-BNST injections of green or red retrobeads (250 nL, Lumafluor) prior to DID to label this population for identification during recordings. For experiments characterizing postsynaptic responses in BNST^CRF^ neurons to PVT glutamate inputs, mice received an intra-PVT stereotaxic injection of AAV5-CamKIIα-hChR2(H134R)-eYFP.WPRE.hGH (200 nL, Penn Vector Core and Addgene) three weeks prior to DID procedures. During recordings, 1 ms 490 nm LED stimulation was used to optically stimulate ChR2+ PVT cell bodies to elicit action potentials to confirm sufficient PVT expression and ChR2 fidelity at 1, 2, 5, 10, 20, and 50 Hz. One 2 ms stimulation every 10 s was used in the BNST to optically-evoke glutamate release from PVT terminals while recording postsynaptic responses (oPSCs) from BNST^CRF^ neurons in voltage-clamp, and 2 ms stimulation at 1, 2, 5, 10, and 20 Hz was used while measuring postsynaptic potentials (oPSPs) in BNST^CRF^ neurons in current-clamp. Polysynaptic and monosynaptic oPSCs were assessed with the bath application of TTX (500 nM) to block all neurotransmission followed by the addition of 4-aminopyridine (100 µM) to reinstate monosynaptic transmission.

Following behavior, a subset of PVT^BNST^ DREADD mice were sacrificed for slice electrophysiology confirmation of the approach using bath application of CNO (10 µM) during current-clamp recordings of DREADD+ PVT cell bodies in the presence of TTX (**Supplementary Fig. 7c**). To provide confirmation of the approach at the BNST^CRF^ neuron population level, CRF-Cre mice received bilateral intra-BNST injections of the calcium sensor GCaMP6s (AAV4-Syn-FLEX-GCaMP6s; UPenn Vector Core; 500 nL) and an intra-PVT injection of the excitatory hM3D DREADD virus (AAV2-CaMKIIα-hM3D(Gq)-mCherry, 200 nL) or control virus (AAV8-CaMKIIα-mCherry; **Supplementary Fig. 7d**). Four weeks later, fresh brain slices were acutely prepared as described for slice electrophysiology recordings above. GCaMP6s was excited with 35% 470 nm LED (CoolLED) at a frequency of 1 Hz for 10 s every min to minimize photobleaching, and videos were acquired at a frame rate of 10 Hz with an optiMOS monochrome camera (QImaging, Surrey, British Columbia, Canada), across the entire experiment including a five min baseline, 10 min bath application of CNO (10 μM), and 10 min washout period. The experimenter maintained objective focus on the BNST z-plane of interest containing several CRF^GCaMP6s^ neurons throughout the experiment. Custom MATLAB code was used to analyze changes in fluorescence intensity in individual CRF^GCaMP6^ neurons (ΔF) compared to background fluorescence within the frame (intensity of entire field of view, F) throughout the video. CRF^GCaMP6s^ cells from CON virus mice were used to quantify the inherent linear decay in fluorescence of GCaMP6s, and fluorescence in hM3D CRF^GCaMP6s^ cell fluorescence was normalized to this decay for statistical analysis.

### Statistical analyses

Statistical analyses were performed in GraphPad Prism, R, and MATLAB. Data for all dependent measures were examined for their distributions in normal and log space, outliers, and equality of variance across groups using Q-Q plots. Electrophysiological properties and synaptic transmission data were lognormally distributed, analyzed using log-transformed values, and presented as raw values on a log2 scale in figures; all other data were normally distributed, analyzed in raw space, and presented on a linear scale in figures. Outliers according to Q-Q plots were excluded (however this was rare and reported). Data are presented as mean ± SEM, and raw data points are included in all figures except those with more than two repeated measures when there were too many raw data points to be clearly represented.

Two-way analysis of variance (ANOVA) and unpaired t-tests were used to evaluate the effects of sex and alcohol on synaptic transmission and excitability data. Repeated measures ANOVAs (RM-ANOVAs) were used to examine the effects of treatment, cycles of binge drinking, different anatomical subregions, etc., within individual animals across experimental groups; mixed effects models were used when one or more matched data point was unavailable for an individual animal. To prevent false positive results and overinterpretation of RM-ANOVAs on behavioral data, sphericity was not assumed and the Geisser and Greenhouse correction of degrees of freedom was employed for the repeated measure. For all ANOVAs, significant effects were further probed with appropriate post hoc paired or unpaired t-tests with Holm-Sidak (H-S) correction for multiple comparisons, and multiplicity-adjusted p values are reported. Differences in proportions between groups were assessed with Fisher’s exact tests, and differences within group from a null hypothesis value were evaluated using one-sample t-tests. Statistical comparisons were always performed with an alpha level of 0.05 and using two-tailed analyses.

## Data availability statement

All data generated or analysed during this study are included in this published article (and its supplementary information files).

## Acknowledgements

We thank Bryan Roth for contributing DREADD viruses and Bradford Lowell for sharing VGLUT2-ires-Cre and CRF-ires-Cre mouse lines bred in our facilities for these studies. We also thank Alexis Kendra, Nakul Yadav, Jared Boyce, and Avanti Shirke for their technical assistance. This research was supported by: NIH grants K99/R00 AA023559 and R01 AA027645, a NARSAD Young Investigator Award, a Stephen and Anna Maria Kellen Foundation Junior Faculty Award (KEP); NIH grants P60 AA011605, R01 AA019454, and U01 AA020911 (TLK); NIH grant T32 DA039080 (OBL); and NIH grant F32 AA025530 (MJS).

## Author Contributions

KEP designed all experiments, OBL, MJS, and KEP collected and analyzed slice electrophysiology data, OBL, JDM, JFD, SAR, JKRI, JAR, and KEP performed and analyzed behavior data, JDM collected and analyzed anatomical tracing data, and JFD and JDM bred mice for all studies. KEP, TRT, and TLK oversaw experiments, KEP and OBL wrote the manuscript, and all authors edited and approved the final version of the manuscript.

## Supplementary Figures

**Supplementary Fig. 1:**
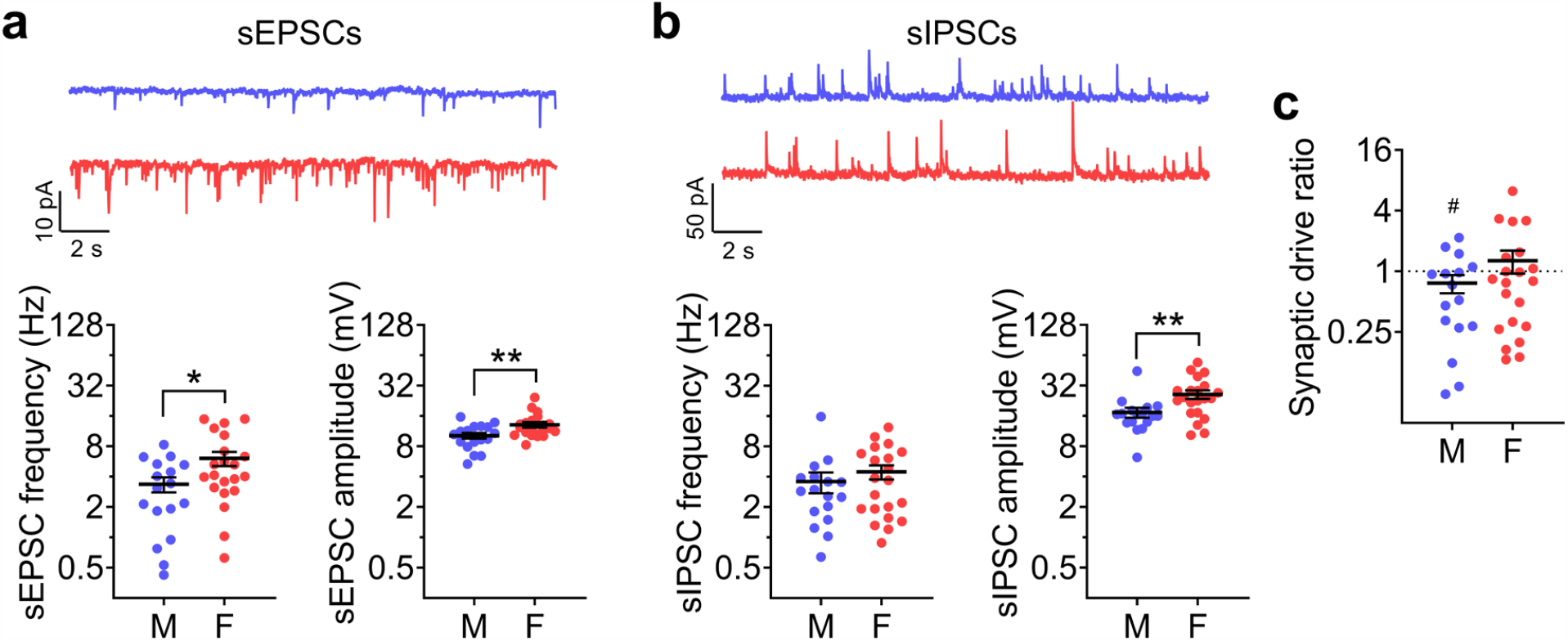
Greater spontaneous glutamate release onto BNST^CRF^ neurons in females than males. Spontaneous excitatory and inhibitory postsynaptic currents (sEPSCs and sIPSCs) in BNST^CRF^ neurons (N’s = 8 M, 17 cells; 8 F, 21 cells). **a**, Top: representative traces of spontaneous excitatory postsynaptic currents (sEPSCs) in BNST^CRF^ neurons of males (blue, above) and females (red, below). Bottom: Quantification showing that sEPSC frequency and amplitude are both higher in females than males (frequency, left: t_36_ = 2.20, \**P* = 0.034; amplitude, right: t_36_ = 2.88, \*\**P* = 0.007). **b**, Top: Representative traces of spontaneous inhibitory postsynaptic currents (sIPSCs) in BNST^CRF^ neurons of males (blue, above) and females (red, below). Bottom: Quantification showing that sIPSC frequency is not different between sexes (left, t_36_ = 0.89, *P* = 0.380) and amplitude is higher in females (right, t_36_ = 3.66, \*\**P* = 0.007). **c**, Synaptic drive ratio, calculated as (sEPSC frequency x amplitude) / (sIPSC frequency x amplitude), in BNST^CRF^ neurons is below 1.0 in males (t_15_ = 2.58, ^#^*P* = 0.021) but not females (t_20_ = 1.35, *P* = 0.191), but there is no difference between males and females (t_35_ = 1.03, *P* = 0.312).

**Supplementary Fig. 2:**
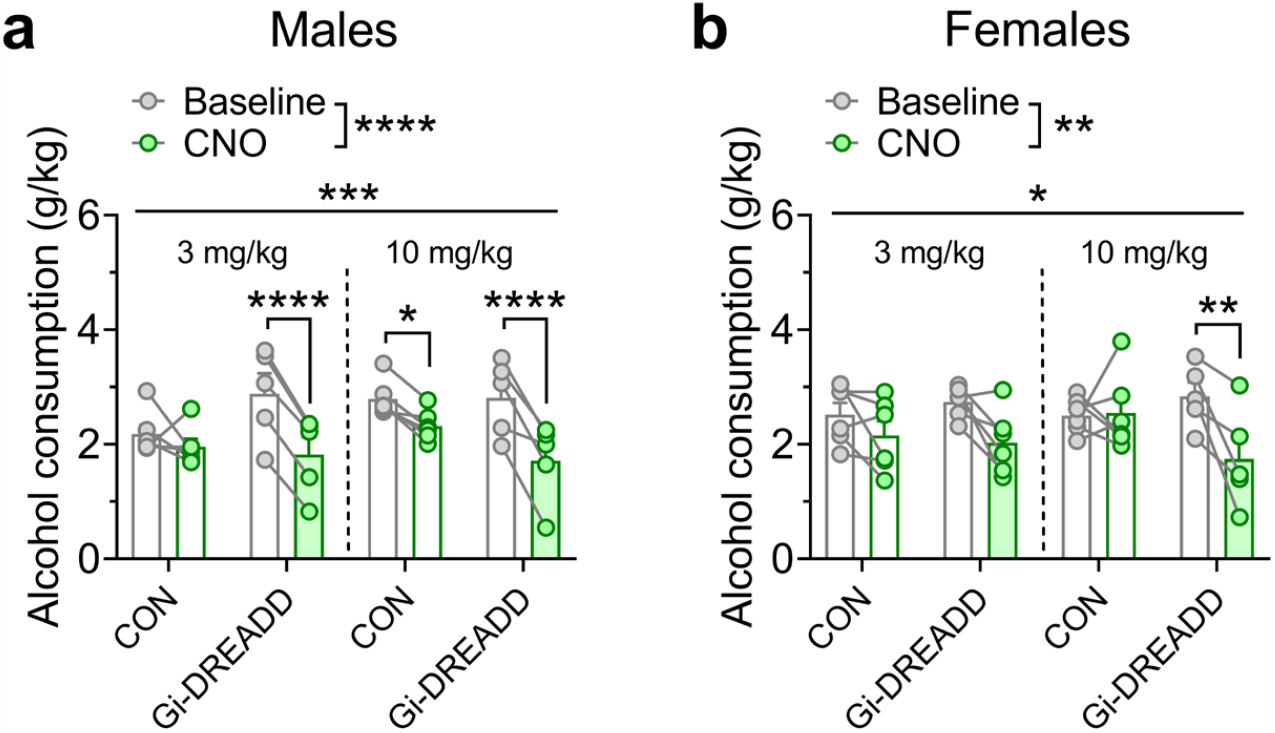
Females require more robust chemogenetic inhibition of BNST^CRF^ neurons to suppress binge alcohol drinking than males. **a**, Binge alcohol drinking in males is inhibited by both moderate (3 mg/kg, i.p.) and high (10 mg/kg, i.p.) dose CNO activation of a Gi-DREADD in BNST^CRF^ neurons (however, there is a nonspecific effect of high dose CNO in CON males). 3xRM-ANOVA: main effect of CNO (F_1,18_ = 68.18, ^****^*P* < 0.0001) and CNO x Gi-DREADD interaction (F_1,18_ = 17.72, ^***^*P* = 0.0005) and no other effects (*P*s > 0.15); post hoc t-tests between vehicle and CNO within group: CON (3): t_18_ = 1.34, *P* = 0.196; Gi-DREADD (3): t_18_ = 5.86, \*\*\*\**P* < 0.0001; CON (10): t_18_ = 2.90, \**P* = 0.019; Gi-DREADD (10): t_18_ = 6.08, \*\*\*\**P* < 0.0001; N’s = 6 CON, 5 Gi-DREADD. **b**, Binge alcohol drinking in females is inhibited by high but not moderate dose CNO activation of a Gi-DREADD in BNST^CRF^ neurons. 3xRM-ANOVA: main effect of CNO (F_1,19_ = 14.18, ^**^*P* = 0.001) and CNO x Gi-DREADD interaction (F_1,19_ = 6.96, ^*^*P* = 0.016) but no other effects (*P*s > 0.15); post hoc t-tests between vehicle and CNO within group: CON (3): t_19_ = 1.34, *P* = 0.355; Gi-DREADD (3): t_18_ = 2.59, *P* = 0.053; CON (10): t_18_ = 0.18, *P* = 0.857; Gi-DREADD (10): t_18_ = 3.63, \*\**P* = 0.007; N’s = 6 CON, 6 Gi-DREADD per sex. (3 mg/kg data are adapted from Pleil et al., 2015 and here represented separately by sex.)

**Supplementary Fig. 3:**
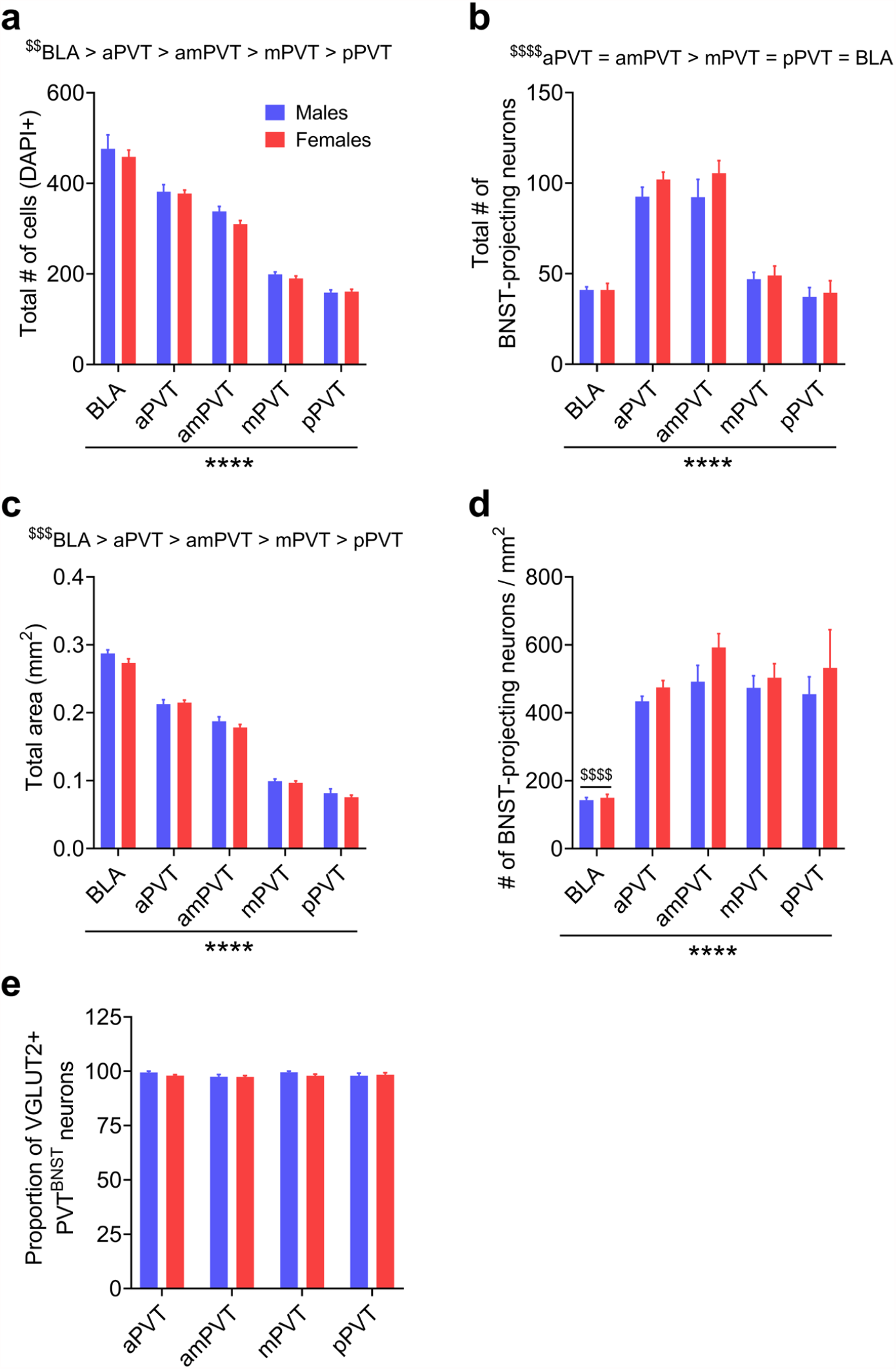
Detailed quantification of glutamatergic inputs to the BNST (related to Fig. 2). **a**, Total number of cells per region/subregion in both sexes, indicated by a DAPI counterstain of nuclei. Mixed-effects model: main effect of subregion (F_4,23_ = 324.0, \*\*\*\**P* < 0.0001) but no effect of sex or interaction (*P*s > 0.25), with BLA > aPVT > amPVT > mPVT > pPVT according to post hoc t-tests (^$$$$^*P*s < 0.0001 for all direct subregion comparisons except mPVT vs. pPVT ^$$^*P* = 0.002). **b**, Total raw number of BNST-projecting neurons across subregions. Mixed-effects model: main effect of subregion (F_4,23_ = 111.1, \*\*\*\**P* < 0.0001) but no effect of sex or interaction (*P*s > 0.30), with aPVT = amPVT > mPVT = pPVT = BLA according to post hoc t-tests (^$$$$^*P*s < 0.0001 for all significantly different subregion comparisons and *P*s > 0.10 for all others). **c**, Total area of each subregion. Mixed-effects model: main effect of subregion (F_4,30_ = 559.9, \*\*\*\**P* < 0.0001) but no effect of sex or interaction (*P*s > 0.05), with BLA > aPVT > amPVT > mPVT > pPVT according to post hoc t-tests (^$$$$^*Ps* < 0.0001 for all direct subregion comparisons except mPVT vs. pPVT ^$$$^*P* = 0.0006). **d**, Number of BNST-projecting neurons normalized to subregion area. Mixed-effects model: main effect of subregion (F_4,23_ = 28.83, \*\*\*\**P* < 0.0001) but no effect of sex or interaction (*P*s > 0.20), with BLA < all PVT subregions (^$$$$^*P*s < 0.0001). **e**, Proportion of PVT^BNST^ neurons that are VGLUT2+ (**Fig. 2e** broken down by A/P subregion, with all sex x subregion proportions > 97.4%. Two-way RM-ANOVA shows no effects or interaction (*P*s > 0.25).

**Supplementary Fig. 4:**
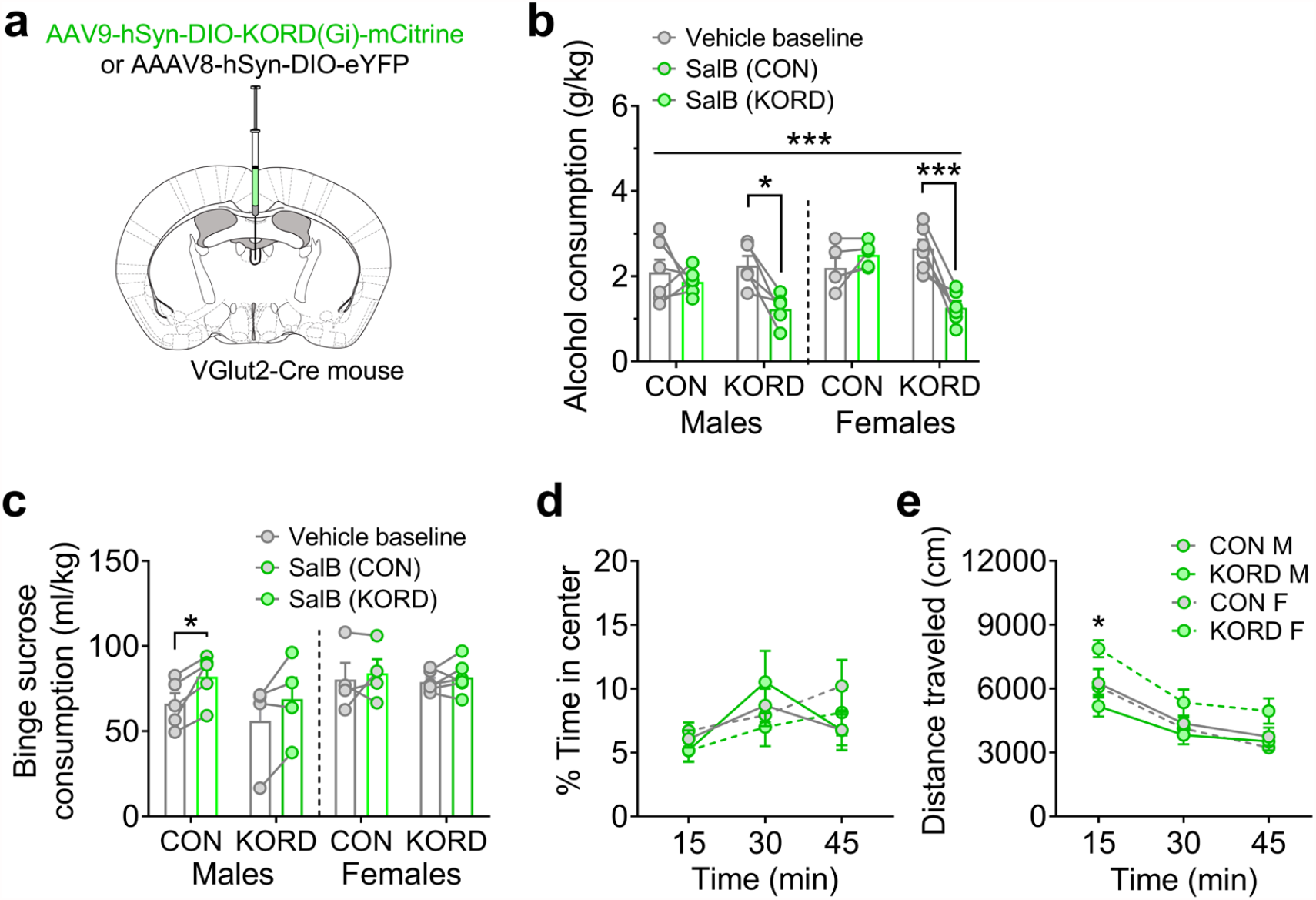
PVT glutamate neurons modulate binge drinking in both sexes. **a**, Schematic of viral strategy to express an inhibitory Gi-KOR DREADD (Gi-KORD) or control virus specifically in PVT^VGLUT2^ neurons. **b**, One-hr binge alcohol consumption during DID at within-cycle baseline including Day 2 vehicle injection (gray) and following Salvinorin B (SalB) injection (17 mg/kg, s.c., green) on Day 4 to activate the KORD (n = 5-6 mice/group). 3xRM-ANOVA: main effects of the KORD (F_1,18_ = 4.92, *P* = 0.037) and SalB (F_1,18_ = 16.17, *P* = 0.0008), and a KORD x SalB interaction (F_1,18_ = 18.51, \*\*\**P* = 0.0004); post hoc paired t-tests show that SalB administration blunts binge drinking in male and female mice with the KORD (*t*_4_ = 3.79, adjusted \**P* = 0.019 and *t*_5_ = 5.29, adjusted \*\*\**P* = 0.0003, respectively) but not their control-virus counterparts (adjusted *P*s > 0.15). **c**, One-hr sucrose consumption during a matched 4% sucrose DID procedure in KORD and control vector-expressing mice (n = 4-6 mice/group, with 0-1 excluded mouse/group with no baseline sucrose consumption). 3xRM-ANOVA: main effect of SalB (F_1,15_ = 11.88, *P* = 0.004) and a SalB x sex interaction (F_1,15_ = 4.79, *P* = 0.045) but no effect of the KORD or sex or other interactions between the three variables (*P*s > 0.05); paired t-tests show that the effect of SalB was driven by an increase in CON males (*t*_4_ = 3.54, adjusted \**P* = 0.024; all others *P* > 0.10). **d-e**, Effects of SalB activation of the Gi-KORD in PVT^VGLUT2^ neurons prior to behavior in the OF. **d**, Percent time in center. 3xRM-ANOVA: main effect of time (F_1.9,32.4_ = 6.05, *P* = 0.007) and a sex x time interaction (F_2,34_ = 3.69, *P* = 0.035) but no effect of the KORD or sex or any other interactions between the variables (*P*s > 0.35); however, post hoc t-tests show no differences between males and females in each 15 min time bin (*P*s > 0.35). **e**, Total distance during the OF test. 3xRM-ANOVA: main effect of time (F_1.7,28.8_ = 116.6, *P* < 0.0001) and a sex x KORD interaction (F_1,17_ = 7.06, *P* = 0.017) but no effect of the KORD or sex or any other interactions between the variables (*P*s > 0.05); post hoc t-tests show a significant difference between CON and KORD females only during the first 15 min time bin (*t*_8_ = 3.28, adjusted \**P* = 0.033; other *P*s > 0.05) and no differences between CON and KORD males (all adjusted *P*s > 0.55).

**Supplementary Fig. 5:**
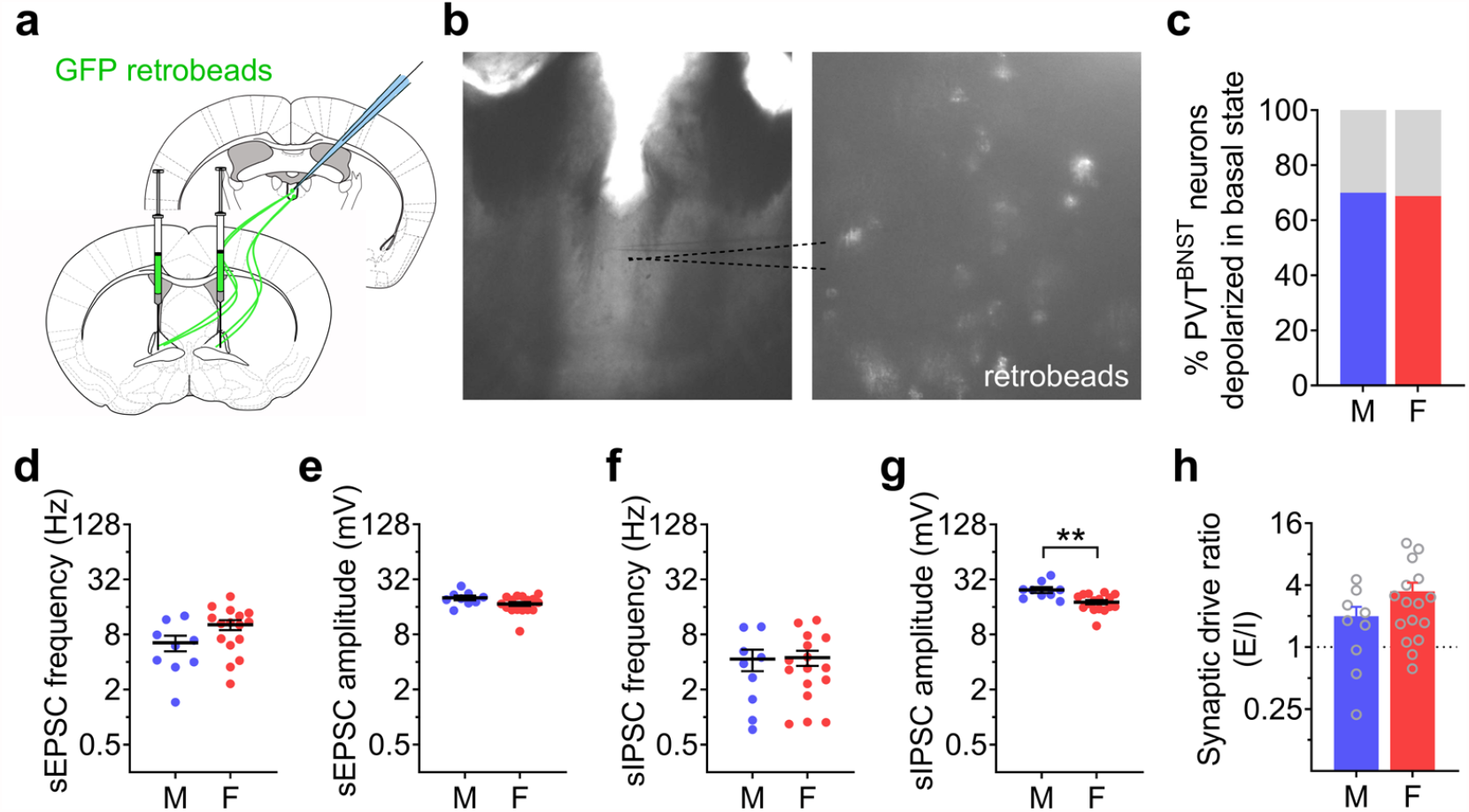
Electrophysiological characterization of PVT^BNST^ neurons. **a**, Retrograde labeling strategy used to identify and perform electrophysiology recordings in BNST-projecting PVT (PVT^BNST^) neurons. **b**, Representative image of a coronal PVT brain slice at 4x (left) containing GFP retrobead-positive cell bodies shown magnified at 40x (right) using monochrome camera on the slice electrophysiology rig. **c**, Percentage of PVT^BNST^ neurons sampled that are active in their basal state is not different between males and females (Fisher’s exact test: *P* > 0.999; N’s = 8 M, 20 cells; 6 F, 19 cells). **d-h**, Synaptic transmission measures from PVT^BNST^ neurons (N’s = 4 M, 9 cells; 5 F, 16 cells; unpaired t-tests). **d**, sEPSC frequency (t_23_ = 1.87, *P* = 0.075). **e**, sEPSC amplitude (t_23_ = 1.86, *P* = 0.075). **f**, sIPSC frequency (t_23_ = 0.20, *P* = 0.840). **g**, sIPSC amplitude (t_23_ = 3.45, \*\**P* = 0.002). **h**, Synaptic drive ratio (t_22_ = 0.79, *P* = 0.438 with Welch’s correction).

**Supplementary Fig. 6:**
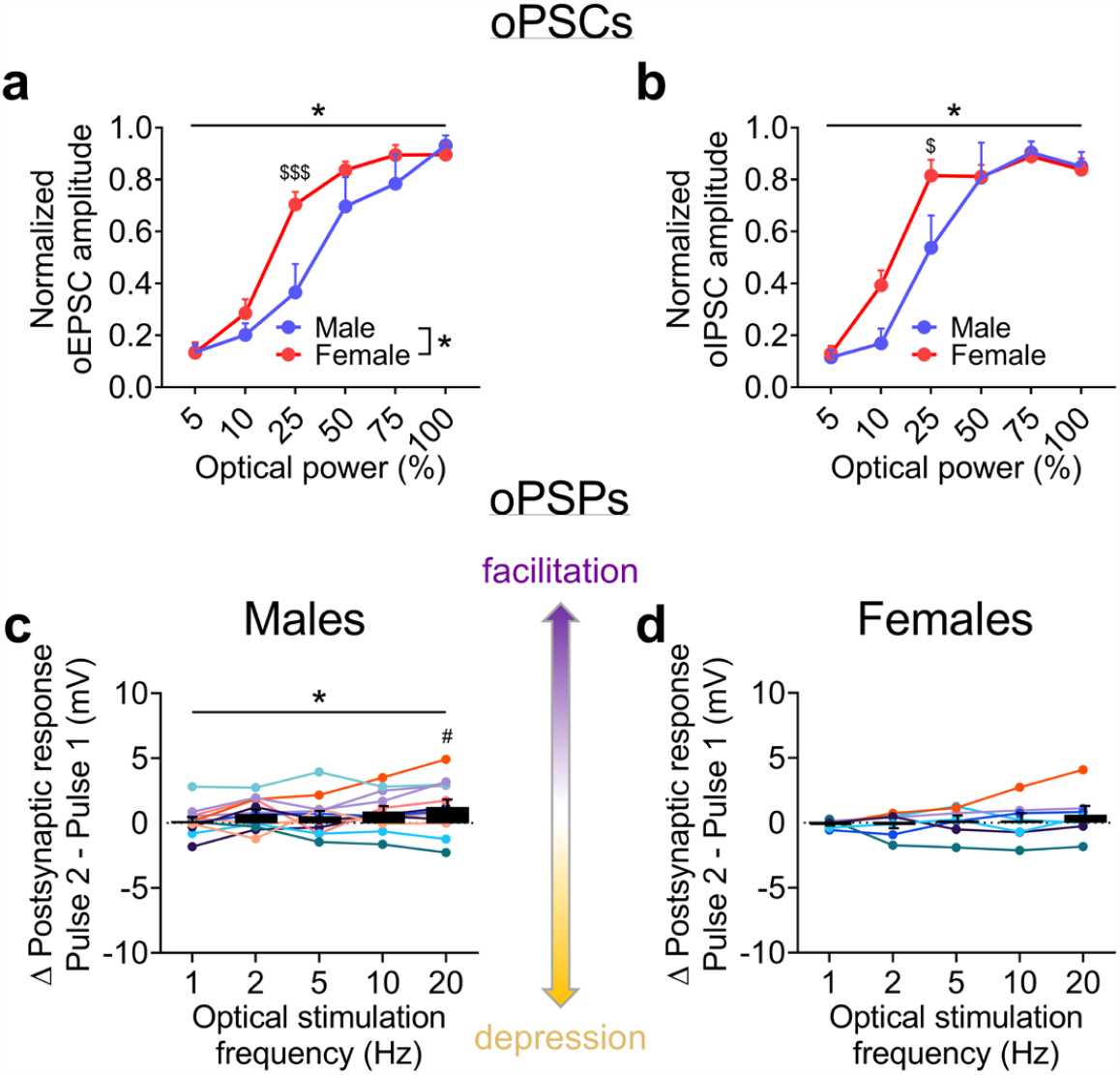
Additional characterization of PVT-BNST^CRF^ synapses (related to Fig. 3). **a-b**, oEPSC (**a**) and oIPSC (**b**) amplitude across LED power (**Fig. 3e**,**f**) normalized to the maximum response within cell shows that females have a left-shifted curves compared to males. oEPSC 2xRM-ANOVA: main effect of sex (F_1,17_ = 6.18, \**P* = 0.024) and power (F_5,85_ = 65.0, *P* < 0.0001, not indicated) and a sex x time interaction (F_5,85_ = 2.69, \**P* = 0.026), with no significant post hoc direct comparisons within LED power (*P*s > 0.10). oIPSC 2xRM-ANOVA: main effect of time (F_5,85_ = 65.0, *P* < 0.0001, not indicated) and a sex x time interaction (F_5,85_ = 2.41, \**P* = 0.043), as well as a trend of sex (F_1,17_ = 4.30, *P* = 0.054), with no significant post hoc direct comparisons within LED power (*P*s > 0.05). **c-d**, Differences between the oPSP elicited from the second and first pulses of LED stimulation across a range of stimulation frequencies within the same BNST^CRF^ neurons during current-clamp recordings, with positive delta values indicating facilitated responses and negative delta values indicating depressed responses (N’s = 7 M, 12 cells; 6 F, 9 cells). **c**, In males, oPSPs from second pulses are larger than first pulses at 20 Hz stimulation but for no other frequencies. 1xRM-ANOVA effect of frequency (F_4,44_ = 2.61, \**P* = 0.025), with one-sample t-tests for delta values at each frequency against the null hypothesis value of 0 showing a significant facilitation at 20 Hz (t_11_ = 2.23, ^#^*P* = 0.048) but no other frequencies (*P*s > 0.05). **d**, In females, oPSPs from first and second pulses of stimulation are similar across all frequencies (1xRM-ANOVA no effect of frequency (F_4,24_ = 0.77, \**P* > 0.55), and no significant one-sample t-tests within frequency (*P*s > 0.35).

**Supplementary Fig. 7:**
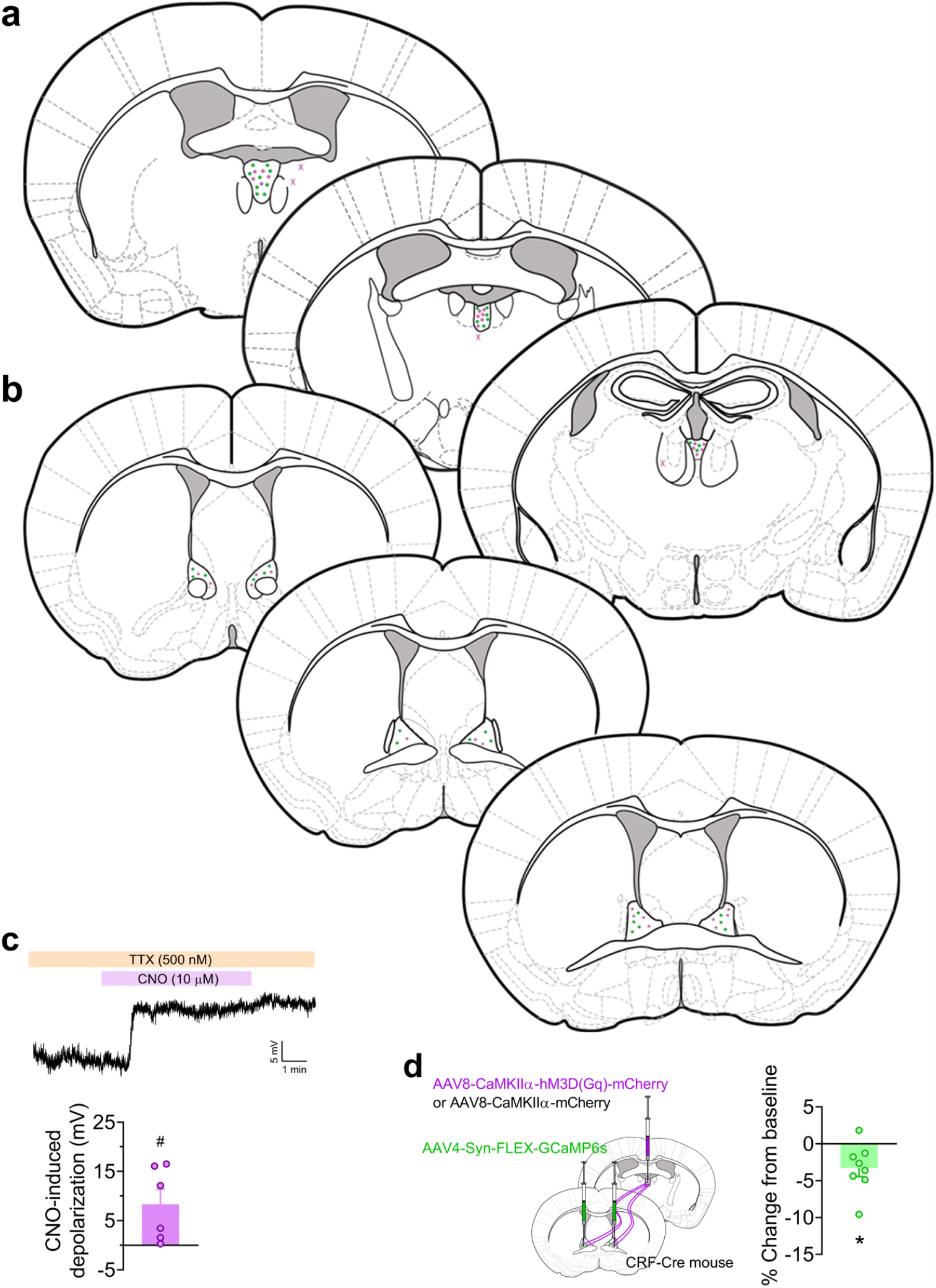
Additional measures of DREADD activation on physiology (related to Fig. 4). **a-b**, Viral injection placements in PVT (**a**) and BNST (**b**) for mice in **Fig. 4**, with hits indicated with circles and misses excluded from analysis in X’s (CONs in green, DREADDs in pink). **c**, Slice electrophysiology in a subset of DREADD mice shows that in the presence of TTX to block action potentials, bath application of CNO (10 μM) depolarizes DREADD+ PVT cell bodies, as shown in the trace above and quantified in the bar graph below (one-sample t-test compared to 0 mV change: t_5_ = 2.75, ^#^*P* = 0.040). **d**, left: Schematic of the strategy to express the calcium biosensor GCaMP6s specifically in BNST^CRF^ neurons and the Gq-DREADD or empty control virus in the PVT for *ex vivo* slice calcium imaging experiment in the BNST during bath application of CNO (10 μM) to activate the Gq-DREADD in PVT terminals. Right: Quantified percent change in fluorescence intensity of GCaMP6s+ in BNST^CRF^ neurons in the presence of CNO from baseline in mice with the Gq-DREADD in the PVT (normalized to background image intensity and standard linear decay of the fluorescence signal observed in neurons from CON virus mice). One-sample t-test: t_7_ = 2.82, \**P* = 0.026. N’s = 1 CON, 6 cells; 2 Gq-DREADD, 8 cells.

**Supplementary Fig. 8:**
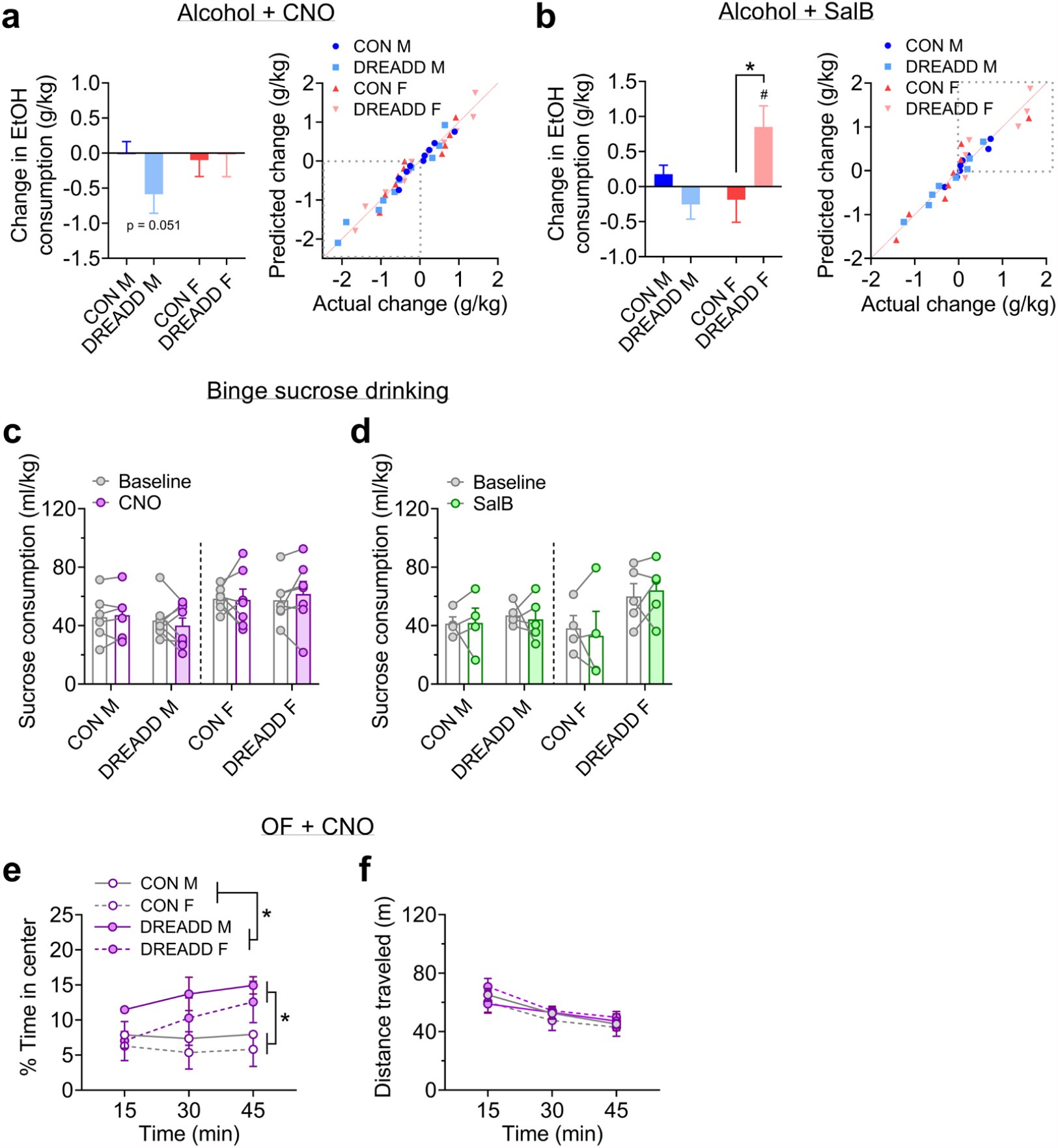
Additional measures of DREADD activation on behavior (related to Fig. 4). **a-b**, Complementary analyses for DID alcohol consumption in **Fig. 4c**,**d. a**, Left: t-tests between CON and DREADD mice within sex comparing the CNO-induced change in drinking show no differences in either sex (*P*s > 0.20). However, one-sample t-tests within each group comparing the change to the null hypothesis delta value of 0 show a trend for a decrease in DREADD males (t_10_ = 2.18, *P* = 0.055) but no change in other groups (*P*s > 0.65), suggesting that activation of the Gq-DREADD in PVT^BNST^ neurons may be sufficient to reduce binge drinking in a subset of DREADD males (see **Fig. 4c**). Right: QQ plots comparing actual change to predicted change in alcohol consumption, demonstrating that all groups’ data come from the same distribution in raw space. The lower left quadrant is outlined with a dashed line to illustrate that it is represented primarily by DREADD M data. **b**, Left: Supporting the results shown in **Fig. 4d**, one-sample t-tests within each group show that only DREADD females have a significant SalB-induced increase in binge drinking (DREADD F: t_5_ = 2.83, ^#^*P* = 0.037; all other *P*s > 0.20). In addition, the change in DREADD F is significantly larger than that in CON F (t_26_ = 2.85, \**P* = 0.017), but this is not the case for males (*P* > 0.20). Right: QQ plots comparing actual change to predicted change in alcohol consumption, demonstrating that all groups’ data come from the same distribution in raw space. The upper right quadrant is outlined with a dashed line to illustrate that it is represented primarily by DREADD F data. **c-d**, DREADD activation does not affect two-hr binge sucrose consumption. **c**, CNO administration. 3xRM-ANOVA: main effect of sex (F_1,24_ = 6.77, *P* = 0.016, not indicated) but no other effects or interactions (*P*s > 0.30). N’s = 6 CON M, 8 DREADD M, 7 CON F, 7 DREADD F. **d**, SalB administration. 3xRM-ANOVA: no effects or interactions (*P*s > 0.05). N’s = 4 CON M, 5 DREADD M, 4 CON F, 5 DREADD F. **e-f**, CNO administration decreases avoidance of the center of the OF in DREADD mice without altering locomotion, confirming the anxiolytic effect in the EPM in **Fig. 4e-g. e**, 3xRM-ANOVA on the percent time spent in the center of the OF shows main effects of DREADD (F_1,7_ = 8.39, \**P* = 0.023) and time (F_1.9,13.5_ = 4.18, *P* = 0.040, not indicated), and a DREADD x time interaction (F_2,14_ = 5.33, \**P* = 0.019). **f**, 3xRM-ANOVA on the distance traveled shows a main effect of time (F_1.5,10.3_ = 56.18, *P* < 0.0001, not indicated), but no other effects or interactions (*P*s > 0.15). N’s = 3 CON M, 3 DREADD M, 3 CON F, 2 DREADD F.

**Supplementary Fig. 9:**
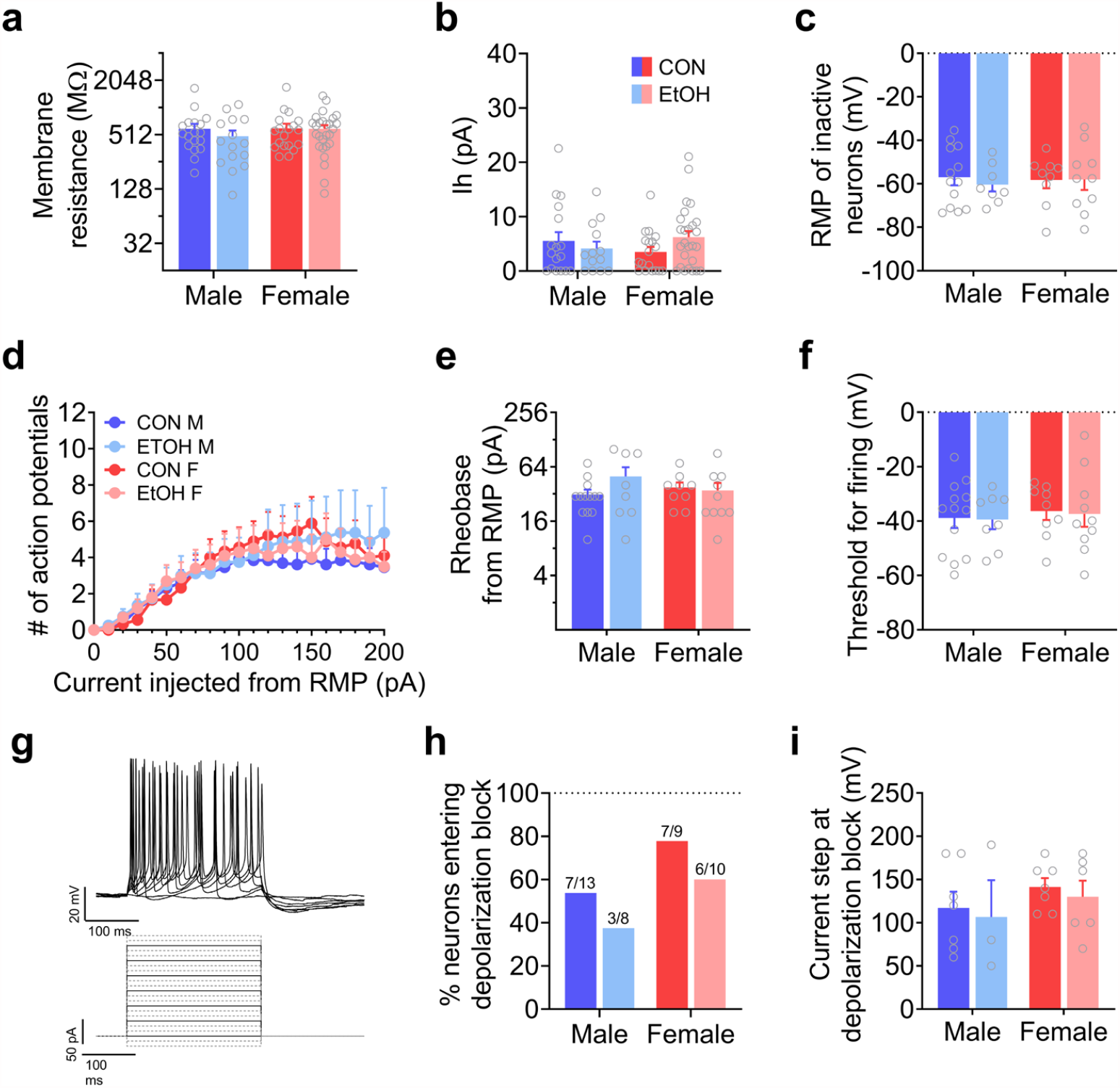
Effects of sex and alcohol on neuronal excitability in BNST^CRF^ neurons at RMP (related to Fig. 5b). **a**, Membrane resistance measured from an I-V plot in voltage clamp configuration with hyperpolarizing steps. 2xANOVA: no effects (*P*s > 0.25). **b**, Hyperpolarization-activated depolarizing current (I_h_) measured in the I-V plot. 2xANOVA: no effects (*P*s > 0.10). **c**, Resting membrane potential (RMP) of neurons that were not displaying a basal state of activity (firing) during gap-free recordings in current-clamp configuration with no current manipulation. 2xANOVA: no effects (*P*s > 0.65). **d-h**, Measures from a V-I plot in current clamp configuration in which increasing 10 pA steps of current were injected directly into basally inactive cells from their RMP (cells identified and represented in **c**) to induce firing, from -20 pA to 200 pA across sweeps. **d**, Number of action potentials generated across current steps (top), with representative traces from one cell (middle) corresponding to solid lines in the stimulus waveform protocol depicted (bottom). 3xRM-ANOVA: main effect of current step (F_20,720_ = 25.57, *P* < 0.0001) but no other effects or interactions (*P*s > 0.60). **e**, Rheobase (minimum amount of current required to elicit an action potential). 2xANOVA: no effects (*P*s > 0.20). **f**, Voltage threshold for firing. 2xANOVA: no effects (*P*s > 0.55). **g**, Representative trace (above) and stimulus protocol (below) for a V-I plot conducted in BNST^CRF^ neurons at RMP. **h-i**, Proportion of neurons that enter depolarization block within the I-V plot is not different between sex or affected by EtOH (Fisher’s exact test *P*s > 0.35) (**h**) and the step at which this occurs is unaffected (**i**; 2xANOVA: no effects (*P*s > 0.25)). The same lack of effects was observed when a ramp current injection protocol was employed (data not shown). N’s = 9 CON M, 18 cells; 6 EtOH M, 15 cells; 9 CON F, 20 cells; 11 EtOH F, 27 cells. Tonically active cells were excluded from analysis for measures in **c-i**.

**Supplementary Fig. 10:**
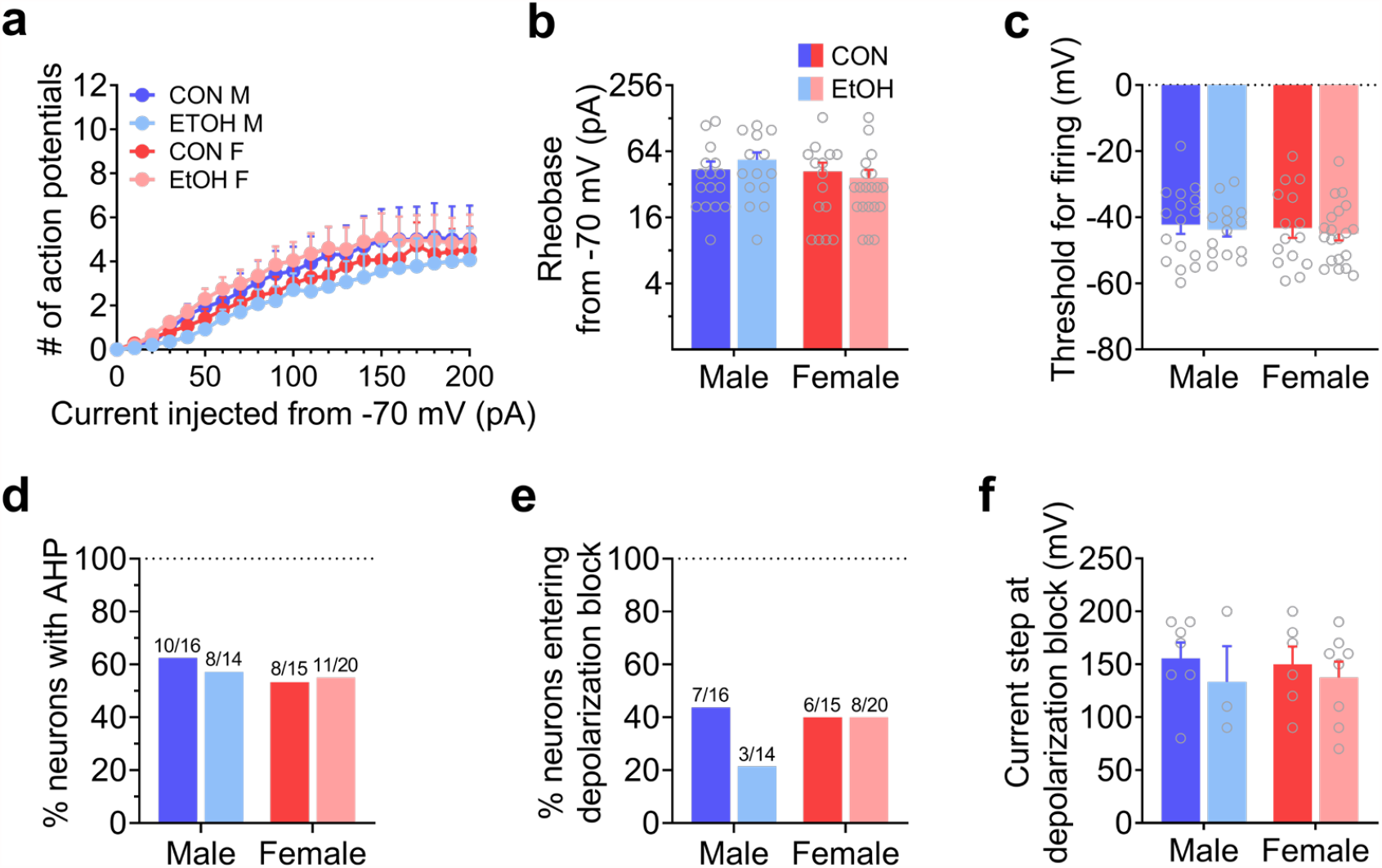
Effects of sex and alcohol on neuronal excitability in BNST^CRF^ neurons when held at a common hyperpolarized membrane potential of -70 mV. **a**, Number of action potentials generated across current steps during a V-I plot starting at a holding potential of -70 mV in all neurons except those displaying firing at this potential (same as **Supplementary Fig. 7d-i** but starting from -70 mV holding potential). 3xRM-ANOVA: main effect of current step (F_20,1220_ = 38.32, *P* < 0.0001) but no other effects or interactions (*P*s > 0.25). **b**, Rheobase. 2xANOVA: no effects (*P*s > 0.10). **c**, Voltage threshold for firing. 2xANOVA: no effects (*P*s > 0.50). **d**, Percentage of neurons displaying an after-hyperpolarization potential (AHP) following action potentials elicited by current injection during the V-I plot is not different between sex or affected by EtOH (Fisher’s exact test *P*s > 0.70). **e-f**, Proportion of neurons that enter depolarization block within the V-I plot is not different between sex or affected by EtOH (Fisher’s exact test *P*s > 0.25). (**e**) and the step at which this occurs is unaffected (**f**; 2xANOVA: no effects (*P*s > 0.35)). The same lack of effects was observed when a ramp current injection protocol was employed (data not shown). N’s are the same as **Supplementary Fig. 7** prior to exclusion based on firing activity at -70 mV.

**Supplementary Fig. 11:**
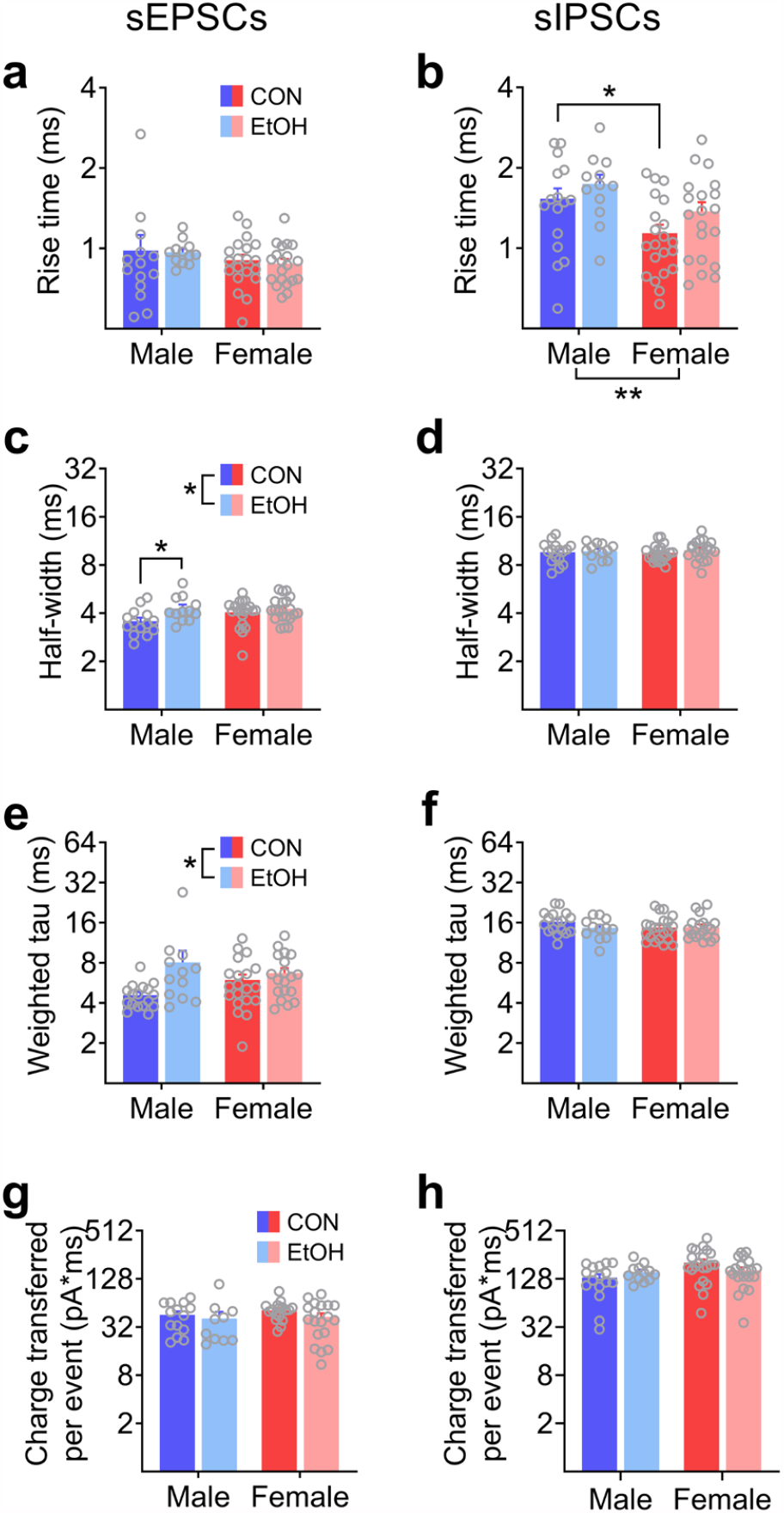
Sex differences and alcohol-induced plasticity in sPSC kinetics in BNST^CRF^ neurons (related to Fig. 5c-h). **a**, sEPSC rise time. 2xANOVA: no effects or interactions (*P*s > 0.30). **b**, sIPSC rise time. 2xANOVA: main effect of sex (F_1,65_ = 9.59, \*\**P* = 0.003) but no other effects (*P*s > 0.05), with a post hoc t-test confirming a sex difference between controls (t_65_ = 2.37, \**P* = 0.021). **c**, sEPSC half-width. 2xANOVA: main effect of alcohol (F_1,62_ = 6.48, \**P =* 0.013) but no other effects (*P*s > 0.15); post hoc t-tests show an effect of alcohol in males (t_62_ = 2.52, \**P* = 0.028) but not females (*P* > 0.35). **d**, sIPSC half-width. 2xANOVA: no effects (*P*s > 0.40). **e**, sEPSC weighted tau. 2xANOVA: main effect of EtOH (F_1,63_ = 6.13, \**P* = 0.016) but no other effects (*P*s > 0.15); post hoc t-tests show that the effect of EtOH was not significant in either sex (*P*s > 0.05). **f**, sIPSC weighted tau. 2xANOVA: no effects (*P*s > 0.20). **g**, sEPSC charge transferred per event, calculated as average area per sEPSC. 2xANOVA: no effects (*P*s > 0.05). **h**, sIPSC charge transferred per event. 2xANOVA: no effects (*P*s > 0.05). For all analyses, cells with an average amplitude of 5 pA or lower after filtering were excluded from analysis—for sEPSCs: 3 CON M, 1 EtOH M, 1 EtOH F; for sIPSCs: 1 CON M, 1 EtOH F. Effects for rise time and half-width in **a-d** were maintained when normalized to the PSC amplitude (not shown).

**Supplementary Fig. 12:**
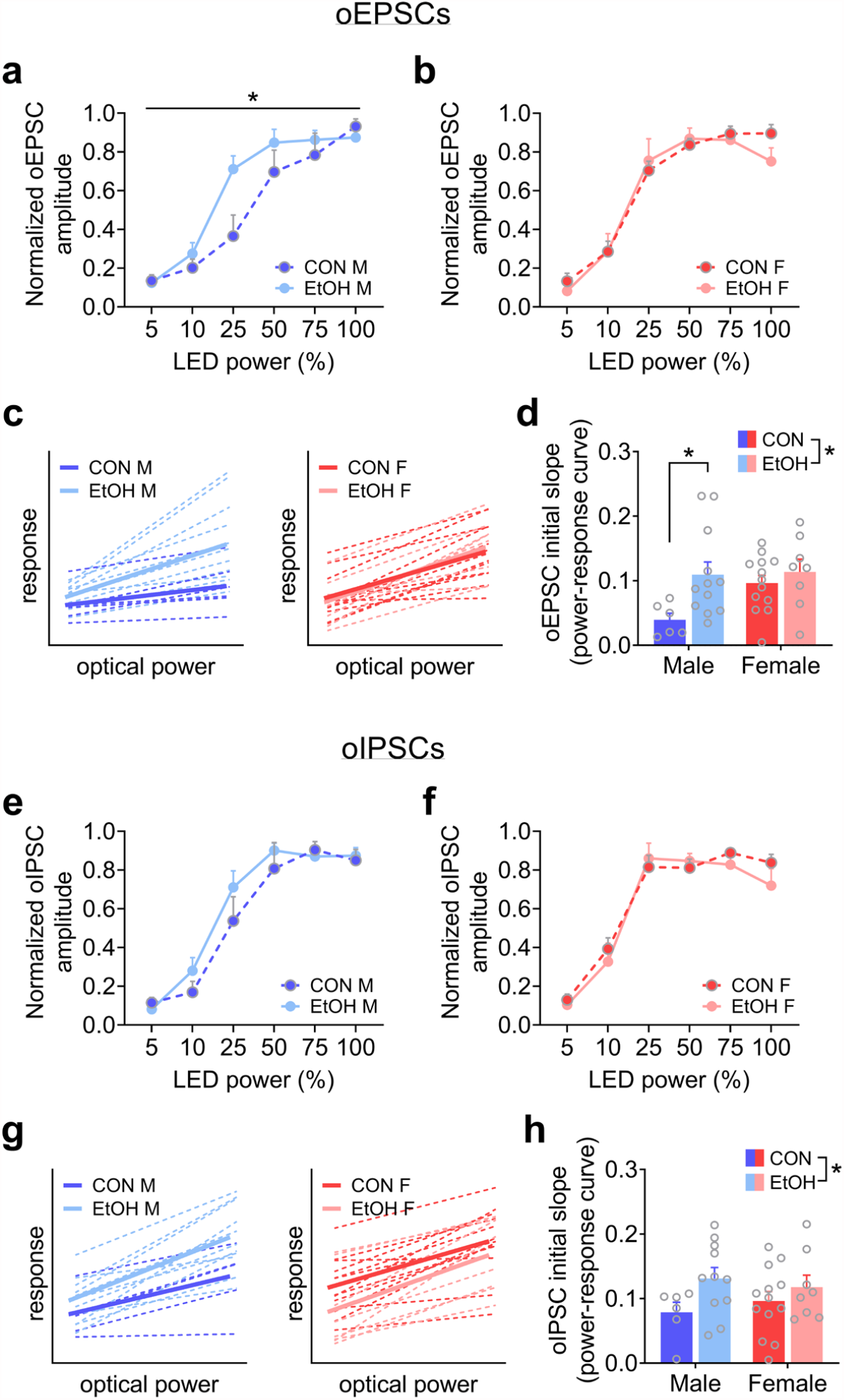
Complementary analyses of PVT-evoked oPSCs (related to Fig. 5i-k). **a-b**, oEPSC amplitude across LED power normalized to the maximum response within cell shows that a history of alcohol drinking results in increased amplitude at lower optical power (leftward shift in the power-response curve) in males (**a**), while it does not affect responses in females (**b**). Males: 2xRM-ANOVA shows a EtOH x time interaction (F_5,80_ = 2.50, \**P* = 0.037) and a main effect of power (F_5,80_ = 51.57, *P* < 0.0001, not indicated) but no main effect of EtOH (*P* > 0.10) and no significant differences in post hoc comparisons within LED power (*P*s > 0.10). Females: 2xRM-ANOVA shows a main effect of power (F_5,95_ = 73.98, *P* < 0.0001, not indicated) but no other effects (*P* > 0.50). **c**, Linear regression for PVT-evoked oEPSCs across 5, 10, and 25% LED power in BNST^CRF^ neurons of males (left) and females (right), with dashed lines showing fits for individual cells and solid lines showing the group average. Regression equations: CON M (Y = 0.03946*X + 1.462), EtOH M (Y = 0.1095*X + 1.438), CON F (Y = 0.09632*X + 1.461), EtOH F (Y = 0.1137*X + 1.209). **d**, Quantification of the oEPSC slope calculated in **c**. 2xANOVA: main effect of EtOH (F_1,35_ = 6.01, \**P* = 0.019) but no effect of sex or interaction (*P*s > 0.05); post hoc t-tests show this effect of EtOH was driven by males (t_35_ = 2.64, \**P* = 0.025) but did not occur in females (*P* > 0.45). **e-f**, oIPSC amplitude across LED power normalized to the maximum response within cell shows no effect of alcohol on PVT-BNST^CRF^ synapses in either sex. **e**, Males: 2xRM-ANOVA shows a main effect of power (F_5,80_ = 57.73, *P* < 0.0001, not indicated) but no other effects (*P* > 0.25). **f**, Females: 2xRM-ANOVA shows a main effect of power (F_5,95_ = 62.69, *P* < 0.0001, not indicated) but no other effects (*P* > 0.35). **g**, Linear regression for PVT-evoked oIPSCs across 5, 10, and 25% LED power in BNST^CRF^ neurons. Regression equations: CON M (Y = 0.07869*X + 1.624), EtOH M (Y = 0.1312*X + 1.926), CON F (Y = 0.09628*X + 2.634), EtOH F (Y = 0.1178*X + 1.523). **h**, Quantification of the oIPSC slope for each BNST^CRF^ neuron calculated in **g**. 2xANOVA: main effect of EtOH (F_1,34_ = 4.42, \**P* = 0.043) but no effect of sex or interaction (*P*s > 0.35); there are no differences in post hoc direct comparisons (*P* > 0.10).

**Supplementary Fig. 13:**
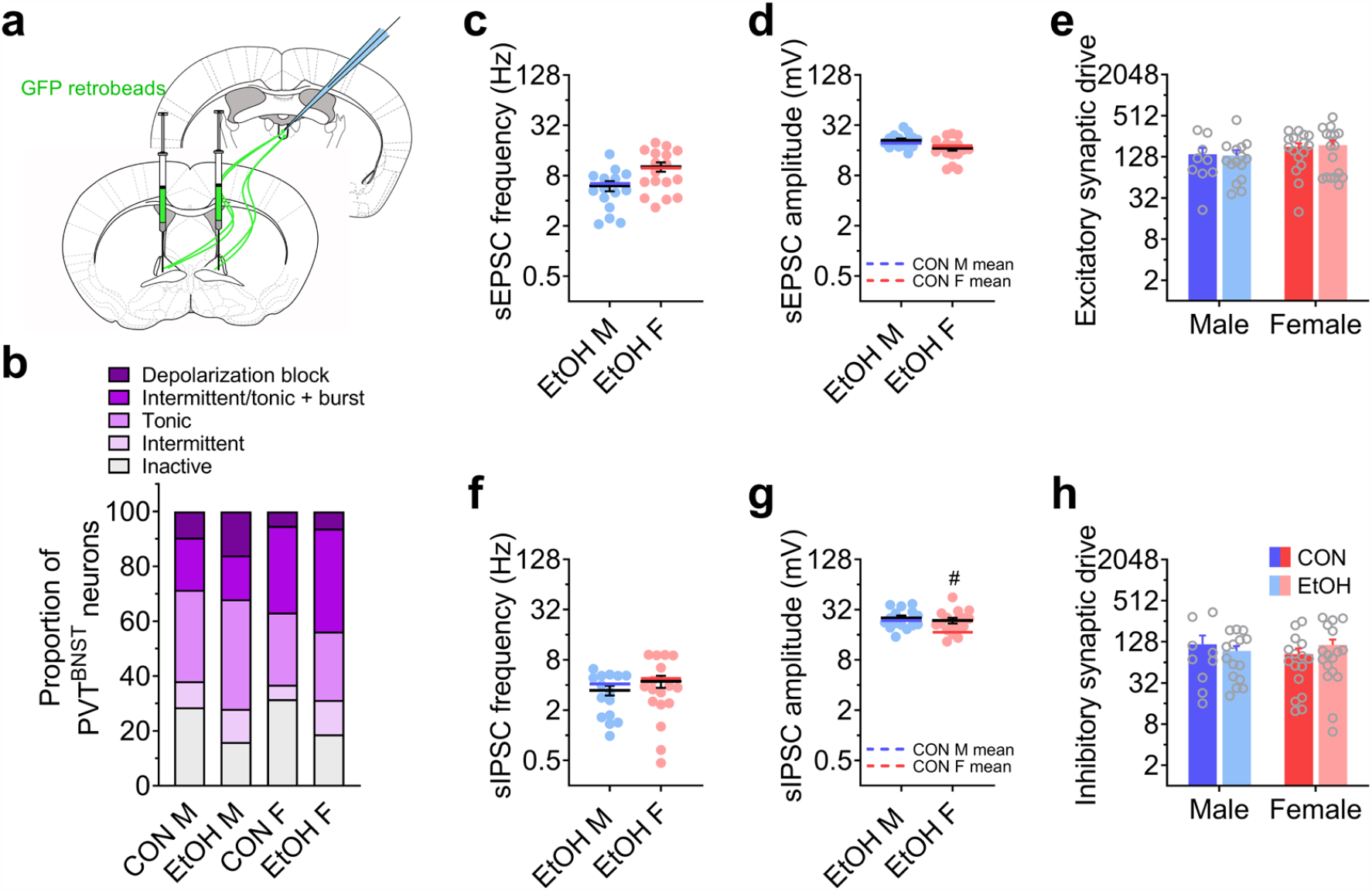
Effects of sex and alcohol on synaptic transmission in PVT^BNST^ neurons. **a**, Schematic showing strategy to label PVT^BNST^ neurons for electrophysiology recordings following 3-cycle DID. **b**, Characterization of the types of neuronal activity in PVT^BNST^ neurons, presented as % sampled population, showing that the population is extremely excitable in all groups without an effect of EtOH in either sex (Fisher’s exact test *P*s > 0.45). N’s = 8 EtOH M, 25 cells; 6 EtOH F, 16 cells. **c-d**, sEPSC frequency (**c**) and amplitude (**d**) in PVT^BNST^ neurons in EtOH M and EtOH F mice (blue and red lines indicating the means in water CON M and F from **Supplementary Fig. 3**). Unpaired t-tests evaluating the effect of EtOH within sex show no effects in either sex for sEPSC frequency (CON M vs. EtOH M: t_22_ = 0.14, *P* = 0.887; CON F vs. EtOH F: t_31_ = 0.13, *P* = 0.990) or amplitude (CON M vs. EtOH M: t_22_ = 0.58, *P* = 0.565; CON F vs. EtOH F: t_31_ = 0.32, *P* = 0.753). **e**, Excitatory synaptic drive, calculated as sEPSC frequency x sEPSC amplitude, within individual neurons. 2xANOVA: no effects (*P*s > 0.55). **f-g**, sIPSC frequency (**f**) and amplitude (**g**) in the same PVT^BNST^ neurons in **c** and **d**. Unpaired t-tests evaluating the effect of EtOH within sex show no effects on sIPSC frequency (CON M vs. EtOH M: t_22_ = 0.111, *P* = 0.912; CON F vs. EtOH F: t_31_ = 0.011, *P* = 0.991), but an effect on sIPSC amplitude in females but not males (CON M vs. EtOH M: t_22_ = 0.175, *P* = 0.863; CON F vs. EtOH F: t_31_ = 2.69, ^#^*P* = 0.011). **h**, Inhibitory synaptic drive within individual neurons. 2xANOVA: no effects (*P*s > 0.70). For **c-h**, N’s = 5 EtOH M, 15 cells; 6 EtOH F, 17 cells.

**Supplementary Fig. 14:**
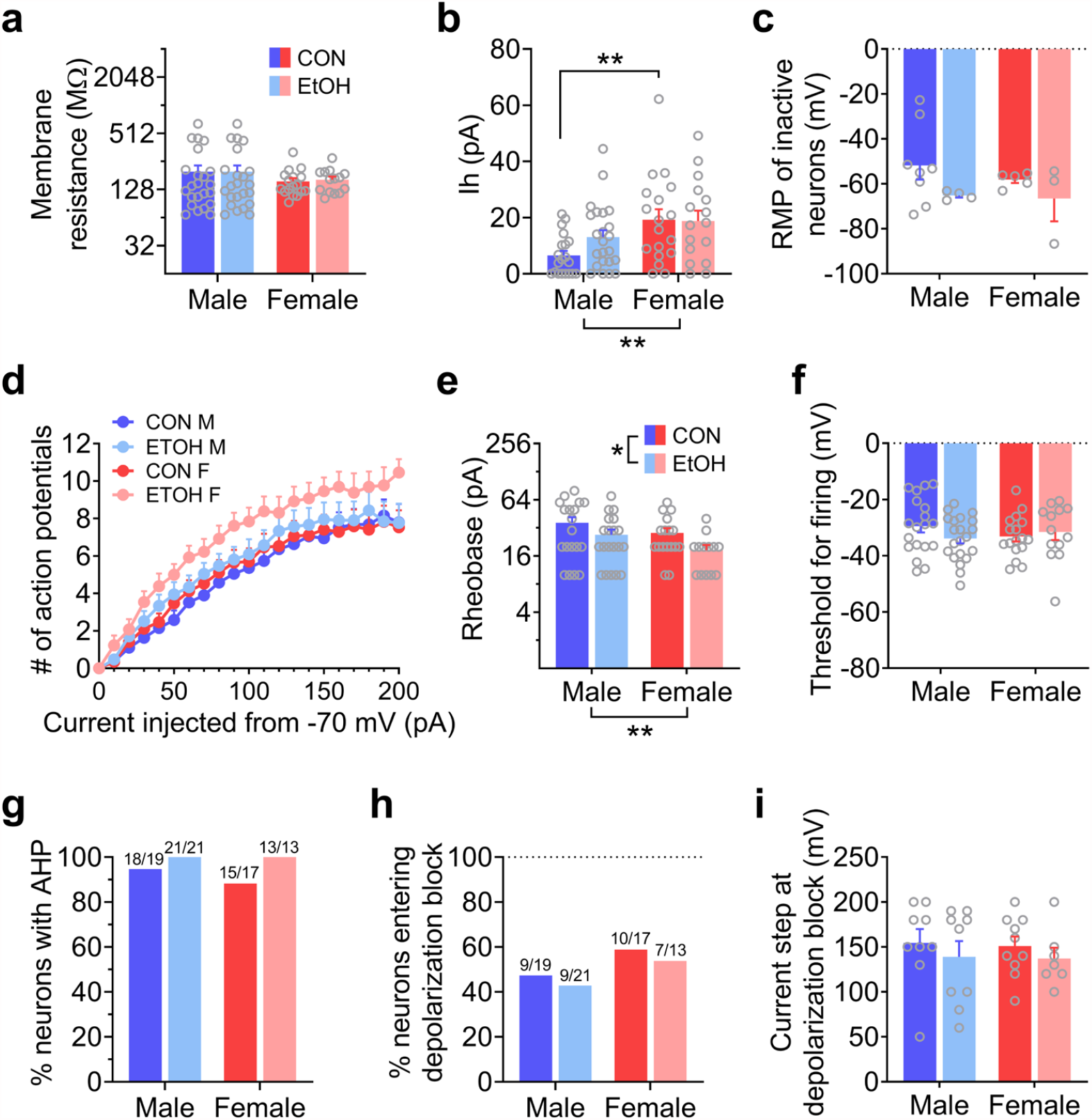
Effects of sex and alcohol on neuronal excitability in PVT^BNST^ neurons. **a**, Membrane resistance measured from an I-V plot in voltage clamp configuration with hyperpolarizing steps. 2xANOVA: no effects (*P*s > 0.80). **b**, Hyperpolarization-activated depolarizing current (Ih) measured in the I-V plot. 2xANOVA: main effect of sex (F_1,73_ = 10.19, \*\**P* = 0.002); a post hoc t-test confirms a sex difference between CON M and F (t_73_ = 3.11, \*\**P* = 0.003). **c**, Resting membrane potential (RMP) of neurons that were not displaying a basal state of activity (firing) during gap-free recordings in current-clamp configuration with no current manipulation. 2xANOVA: no effects (*P*s > 0.10). **d-f**, Measures from a V-I plot in current clamp configuration in which increasing 10 pA steps of current were injected, starting at a holding potential of -70 mV, in all neurons except those displaying firing at this potential. N’s = 8 CON M, 21 cells; 8 EtOH M, 25 cells; 6 CON F, 19 cells; 6 EtOH F, 16 cells. **d**, Number of action potentials generated across current steps. 3xRM-ANOVA: main effect of current step (F_2.5, 164_ = 158.8, *P* < 0.0001) and trend of EtOH (F_1,66_ = 3.87, *P* = 0.054) but no other effects (*P*s > 0.15). **e**, Rheobase. 2xANOVA: main effect of EtOH (F_1,65_ = 5.32, \**P* = 0.024) and main effect of sex (F_1,65_ = 4.03, \**P* = 0.049) but no interaction (*P* > 0.95), with post hoc t-tests showing the effect of alcohol was not driven by one sex (*P*s > 0.15). **f**, Voltage threshold for firing. 2xANOVA: no effects (*P*s > 0.10). **g**, Proportion of neurons displaying an after-hyperpolarization potential (AHP) following spontaneous action potentials is high in all groups and not different between sex or affected by EtOH (Fisher’s exact test *P*s > 0.45; N’s = 8 CON M, 19 cells; 6 CON F, 17 cells; 8 EtOH M, 19 cells; 6 EtOH F, 13 cells). **h-i**, Proportion of neurons that enter depolarization block within the V-I plot is not different between sex or affected by EtOH (Fisher’s exact test *P*s > 0.50) (**h**) and the step at which this occurs is unaffected (**i**; 2xANOVA: no effects (*P*s > 0.30)).

